# A long non-coding RNA at the *cortex* locus controls adaptive colouration in butterflies

**DOI:** 10.1101/2024.02.09.579710

**Authors:** Luca Livraghi, Joseph J. Hanly, Elizabeth Evans, Charlotte J. Wright, Ling S. Loh, Anyi Mazo-Vargas, Kiana Kamrava, Alexander Carter, Eva S.M. van der Heijden, Robert D. Reed, Riccardo Papa, Chris D. Jiggins, Arnaud Martin

## Abstract

Evolutionary variation in the wing pigmentation of butterflies and moths offers striking examples of adaptation by crypsis and mimicry. The *cortex* locus has been independently mapped as the locus controlling colour polymorphisms in 14 lepidopteran species, suggesting it acts as a genomic hotspot for the diversification of wing patterns, but functional validation through protein-coding knockouts has proven difficult to obtain. Our study unveils the role of a novel long non-coding RNA (lncRNA) which we name *ivory*, transcribed from the *cortex* locus, in modulating colour patterning in butterflies. Strikingly, *ivory* expression prefigures most melanic patterns during pupal development, suggesting an early developmental role in specifying scale identity. To test this, we generated CRISPR mosaic knock-outs in five nymphalid butterfly species and show that *ivory* mutagenesis yields transformations of dark pigmented scales into white or light-coloured scales. Genotyping of *Vanessa cardui* germline mutants associates these phenotypes to small on-target deletions at the conserved first exon of *ivory*. In contrast, *cortex* germline mutant butterflies with confirmed null alleles lack any wing phenotype, and exclude a colour patterning role for this adjacent gene. Overall, these results show that a lncRNA acts as a master switch of colour pattern specification, and played key roles in the adaptive diversification of colour patterns in butterflies.

**Significance statement:** Deciphering the genetic underpinnings of adaptive variation is fundamental for a comprehensive understanding of evolutionary processes. Long non-coding RNAs (lncRNAs) represent an emerging category of genetic modulators within the genome, yet they have been overlooked as a source of phenotypic diversity. In this study, we unveil the pivotal role of a lncRNA in orchestrating colour transitions between dark and light patterns during butterfly wing development. Remarkably, this lncRNA gene is nested within the *cortex* locus, a genetic region known to control multiple cases of adaptive variation in butterflies and moths, including iconic examples of natural selection. These findings highlight the significant influence of lncRNAs in developmental regulation, and also underscore their potential as key genetic players in the evolutionary process itself.

## Introduction

The *cortex* locus represents a remarkable example of parallel evolution in butterflies and moths: the same genomic region has repeatedly evolved adaptive alleles that drive phenotypic variation within many diverse lineages. In particular, it underlies colour variation involved in industrial melanism in geometrid moths *Biston, Phigalia* and *Odontopera* (1, 2), adaptive mimicry in *Heliconius* butterflies (3–7) and *Papilio clytia* swallowtails (8), crypsis in leaf-mimicking *Kallima* butterflies (9), seasonal polyphenism in *Junonia* butterflies (10), female-limited melanism in *Pieris napi* (11) and pigmentation mutants in *Bombyx* silkworms (12).

In naturally occuring *Heliconius* hybrid zones, association mapping has implicated discrete, modular non-coding intervals centred around *cortex*, suggesting differences in *cis*-regulatory elements (CREs) are the causal variants driving colour pattern evolution (4, 6, 13, 14). In a captive bred population of *Heliconius melpomene*, a large spontaneous deletion line dubbed *ivory* (here named *ivory*^Δ*78k*^), yields butterflies with depigmented scales in the homozygous state, and involves a 78 kb structural variant that does not contain the protein coding region of *cortex* (15). To date, few hints as to how this locus modulates colouration have been gleaned. CRISPR knock-outs of *cortex* have resulted in rare, small clones with depigmented scale states (6, 10, 16). However, *cortex* is expressed in all epithelial and scale cell precursors of the butterfly wing during pupal development, seemingly with little correlation with adult colour patterns (6), and its molecular function in meiosis-specific cell cycle regulation in *Drosophila* (17) makes it difficult to tie to pigmentation phenotypes. As such, whether *cortex* has a colour patterning role remains an open question, and alternative mechanisms may be reconsidered. For example, prevailing genome annotations often neglect the presence of a class of non-coding transcripts, called long non-coding RNAs (lncRNAs), thus making them easy to miss during genotype-phenotype studies. While a majority of eukaryotic lncRNAs are considered to be transcriptional noise (18), a fraction of them have been described as genuine regulators of gene expression (19, 20). At the *cortex* locus, tiling arrays have revealed that colour morphs of *H. melpomene* show differential transcription of unannotated sequences (4), hinting at the presence of putative lncRNAs within the *ivory*^Δ*78k*^ interval. Nevertheless, assessing the role of lncRNAs in colour patterning would require proper studies of their expression, function, and evolutionary conservation.

Here, the caveats and ambiguities about the role of *cortex* in scale fate specification motivated a closer examination of the genetic elements included in the *ivory* deletion in *Heliconius*, and by extension, of the mechanism behind the various phenotypes previously associated to the *cortex* locus. In parallel to a companion paper by Fandino *et al.* (*in submission*), we find in multiple species that a lncRNA encoding gene, that overlaps with the *ivory*^Δ*78k*^ deletion, is a potent modulator of melanic scale identities. As lncRNAs are being discovered and annotated at an increasing rate in genome databases, the range of their biological functions remain a broadly uncharted territory (19, 21). Importantly, the *ivory* lncRNA shows a conserved function in colour scale specification, while having also evolved divergent expression patterns that precisely delineate pattern information in multiple species. We discuss how these findings not only join a rich literature on the emerging roles of lncRNA in development and gene regulation (22–24), but also imply a direct role in the elaboration of adaptive phenotypic variation.

## Results

### A previously hidden genetic element is expressed at a hotspot locus

In addition to the *ivory^Δ78k^* deletion isolated in a captive stock of *H. melpomene*, several colour polymorphisms encountered in natural hybrid zones have been previously mapped to the *cortex* locus in both *H. melpomene* and *H. erato* (**Fig. 1A-B**). In *H. melpomene*, the causal alleles driving the presence-absence of yellow hindwing bars segregate into two regulatory regions, located 5’ and 3’ of *cortex,* that respectively control the ventral and dorsal wing surface (6, 25, 26). In *H. erato*, the same trait variation is controlled by distinct alleles mapping broadly over *cortex* or its 5’ upstream region, respectively specific to the Western Andes and to Panama (13, 25, 27).

**Figure 1.**
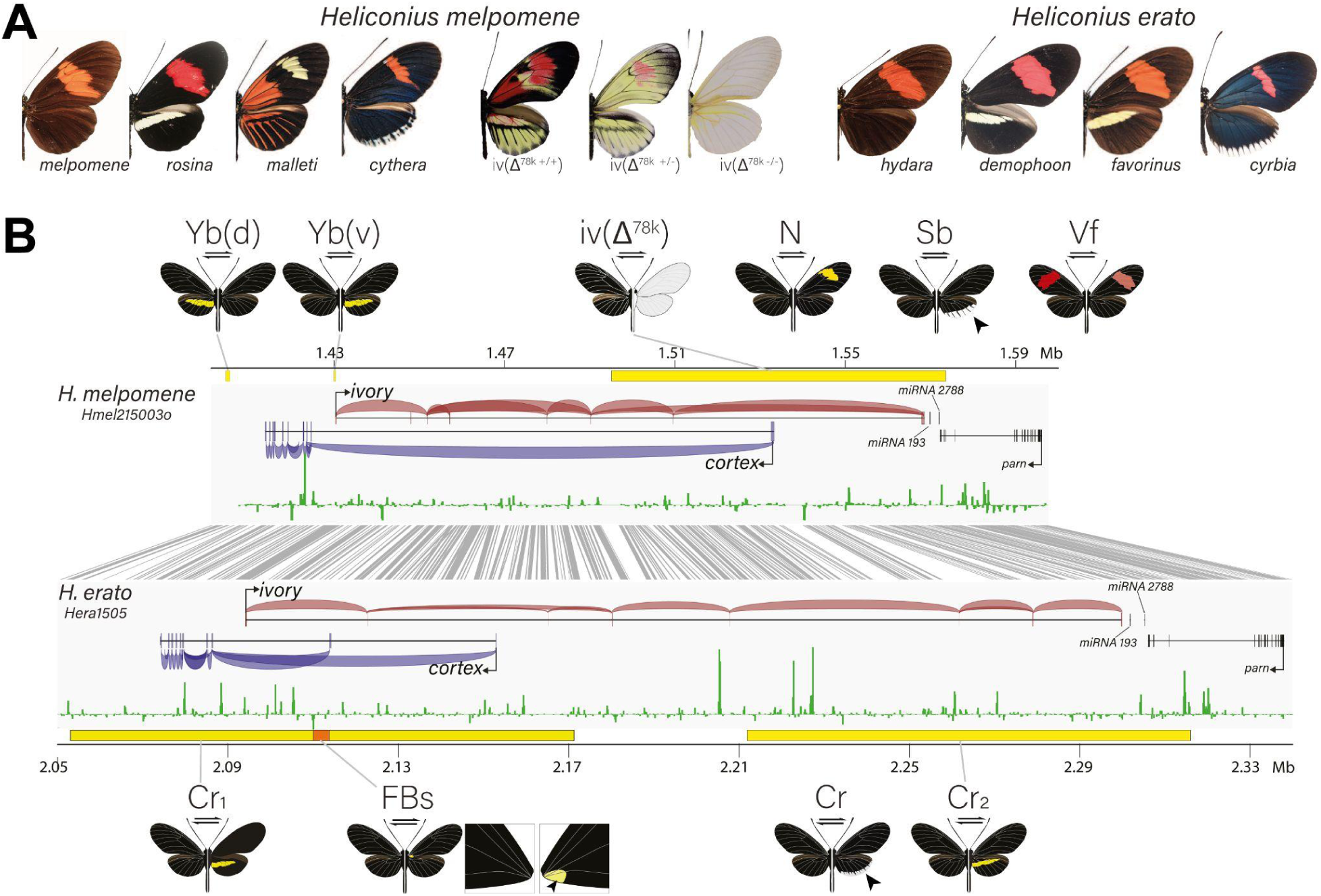
Expression and CRISPR mKOs of the *ivory* lncRNA reveal colour patterning roles. **(A)** Examples of *H. melpomene* and *H. erato* morphs exhibiting colour pattern polymorphisms associated with the *cortex* locus, alongside representative phenotypes of the *ivory*^Δ78k^ deletion. **(B)** Summary of the association intervals previously mapped at the *cortex* locus for *H. melpomene* and *H. erato* (6, 13, 15, 30). For *H. melpomene*: *Yb(d)* dorsal yellow bar (*Hmel215003o*: 1,403,500-1,405,500), *Yb(v)* ventral yellow bar (1,429,500-1,430,500), and *ivory^Δ78k^*deletion (1,494,903-1,573,176). For *H. erato*: *Cr1* Peruvian (West of Andes) yellow bar (*Hera1505*: 2,053,037–2,171,230), *Cr2* Panamanian (East of Andes) yellow bar (2,211,881–2,315,926), and *FBs* forewing base spot (2110000-2113800). Current mapping intervals for *H. melpomene* loci *N*, *Sb* and *Vf*, and the *H. erato Cr* locus are too large to indicate on the figure (31–34). Combined RNA-seq read junction events are shown for both *ivory* (red, positive strand) and *cortex* (blue, negative strand), with annotations for the adjacent *miRNAs 193, 2788* (35) and for the gene *parn*. ATAC-seq tracks from normalised average read depth from 36 h pupal hindwings, from which normalised caterpillar head ATAC-seq depth data has been subtracted, is shown in green. Grey lines indicate aligned regions > 250 bp between *H. melpomene* and *H. erato*.

To further explore the presence of genetic features in this genomic region, we re-analysed RNA-seq data from pupal wings of *H. melpomene* and *H. erato* (28) and identified a polyadenylated, alternatively-spliced lncRNA that is transcribed upstream of *cortex*, hereafter called *ivory* (**Figs. S1-3**). Both reference-based RNAseq alignments and *de novo* transcriptome assemblies reveal that *ivory* encodes a ∼1.3 kb transcript, containing up to eight exons spanning over 138 kb in *H. melpomene* and 205 kb in *H. erato* (**Fig. 1B**). During fifth instar larval wing development, *cortex* transcription initiates from a distal promoter, while *ivory* expression is absent (**Fig. S1-S2**). From 36 hr after pupa formation, or ∼20% of pupal development, *cortex* expression significantly decreases and *ivory* is detected and persists up until at least 60 hr after pupa formation. In *H. erato*, we identified an additional lncRNA transcribed only in fifth instar caterpillars, sharing several antisense exons with *cortex* (**Fig. S1**). In addition, ATAC-seq open-chromatin profiling from both species (29) showed that the *ivory* first exon is immediately 3’ of a region of differential accessibility specific to pupal wing tissues, consistent with a promoter function (**Figs. 1B and S1**).

### The *ivory* lncRNA is necessary for melanic scale development

In order to investigate potential functional roles of the *ivory* lncRNA, we next used CRISPR/Cas9 mutagenesis targeting the *ivory* promoter and first exon in order to test its putative function in wing patterning. These mosaic knock-outs, conducted in *H. erato, H. charithonia* and *Vanessa cardui,* generated butterflies with pronounced shifts in pigmentation, consisting of scale transformations from melanic to yellow-white states (**Fig. 2 and Fig. S4-S5**), phenocopying the effects of the *ivory^Δ78k^* mutant in *H. melpomene* (15). The high penetrance of this phenotype (in 57% of adults), as well as the large size of mutant clones, often spanning entire individuals that emerged healthily (**Fig. 2 and Table S1-S2**), suggests a low pleiotropy of the *ivory* CRISPR phenotypes, with no detectable effects other than in epithelial scales throughout the adult wing and body. Moreover, *in situ* hybridisations against the first exon of *ivory* showed that the *ivory* lncRNA is expressed in perfect correlation with melanic scales during pupal wing development, an association that was previously not found for the *cortex* mRNA or protein (Livraghi et al. 2021). Together these data strongly suggest that *ivory* is the causative locus generating pattern divergence in nymphalid butterflies, rather than *cortex* as previously thought (4, 6, 9).

**Figure 2.**
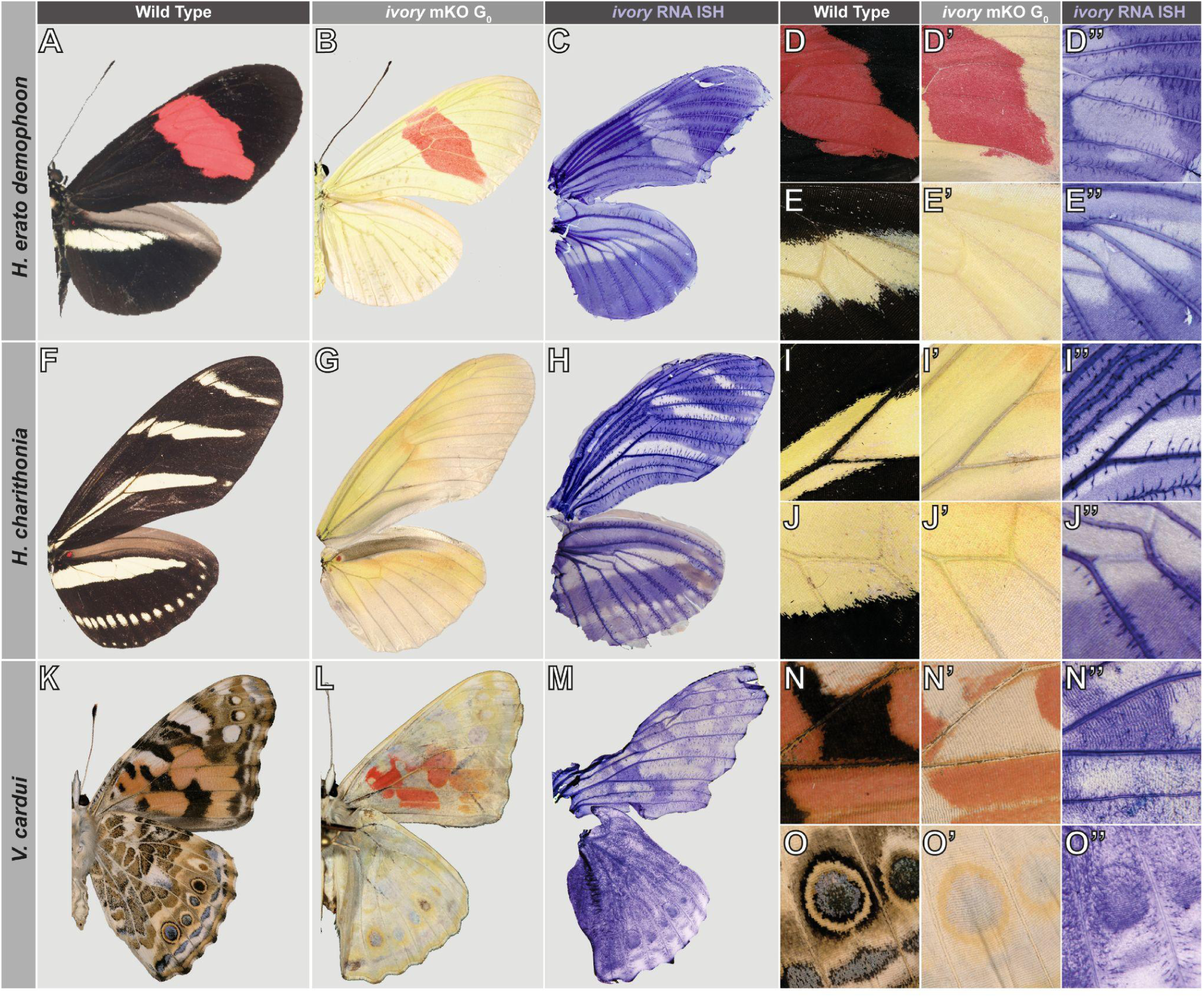
Expression of *ivory* associates with melanic scales in nymphalid butterflies. **(A-C)** Expression of *ivory* in 30% pupal wing tissue of *H. erato demophoon* prefigures adult melanic scales, and mKO results in a complete lack of black patterns. Expression is absent in presumptive red **(D-D’’)** and yellow **(E-E’’)** regions. **(F-J’’)** Expression and function are conserved in *H. charithonia*, strongly marking future black scales, resulting in yellow/white states in the mKOs. **(K-M)** Function is conserved outside *Heliconius*, with *V. cardui* displaying similar loss of melanic scales in the mKOs and a strong correlation between *ivory* expression and adult black patterns. **(N-N’’)** Expression of *ivory* is absent from ventral forewing orange-red ommochrome scales, as well as from **(O-O’’)** white margin patterns and yellow eyespot rings.

### Phenotypic effects are specific to the *ivory* lncRNA

CRISPR-induced mutant clones may include deletions that are larger than intended, particularly in G_0_ injected individuals, where such effects are difficult to genotype. To overcome the limitations of mosaic knock-outs, we thus sought to generate *ivory* KO lines in the Painted Lady butterfly *V. cardui*, a species amenable to mass-rearing in laboratory conditions. We generated G_1_ and G_2_ *V. cardui ivory* mutants by crossing G_0_ individuals displaying mosaic *ivory* mutations, followed by G_1_ pooled matings. This generated viable mutants displaying marked reductions of melanin on all scales, including in the thorax, head, abdomen and antennae (**Fig. 3D-F and Fig. S5-S6**). Tyrosine and Tryptophan radiolabelling experiments – melanin and ommochrome precursors respectively – indicate a strong correspondence between *ivory* KOs and melanin-containing scales (**Fig.3D,E and Fig. S7-S9**). Whole-genome sequencing of G_1_ and G_2_ individuals revealed that compound heterozygosity at the targeted site, typically consisting of two deletion alleles of unequal size, was associated with *ivory* phenotypes of variable expressivity (**Fig. S10**). For example, we established that a bi-allelic deletion spanning 82/97 bp across the lncRNA first exon is sufficient to generate a strong *ivory* phenotype (**Fig. 3F**). These associations between *ivory* exonic deletions and the colouration phenotype, together with the spatial expression of the lncRNA in association with the affected patterns, show that expression of the *ivory* lncRNA positively regulates melanic scale differentiation during pupal wing development.

**Figure 3.**
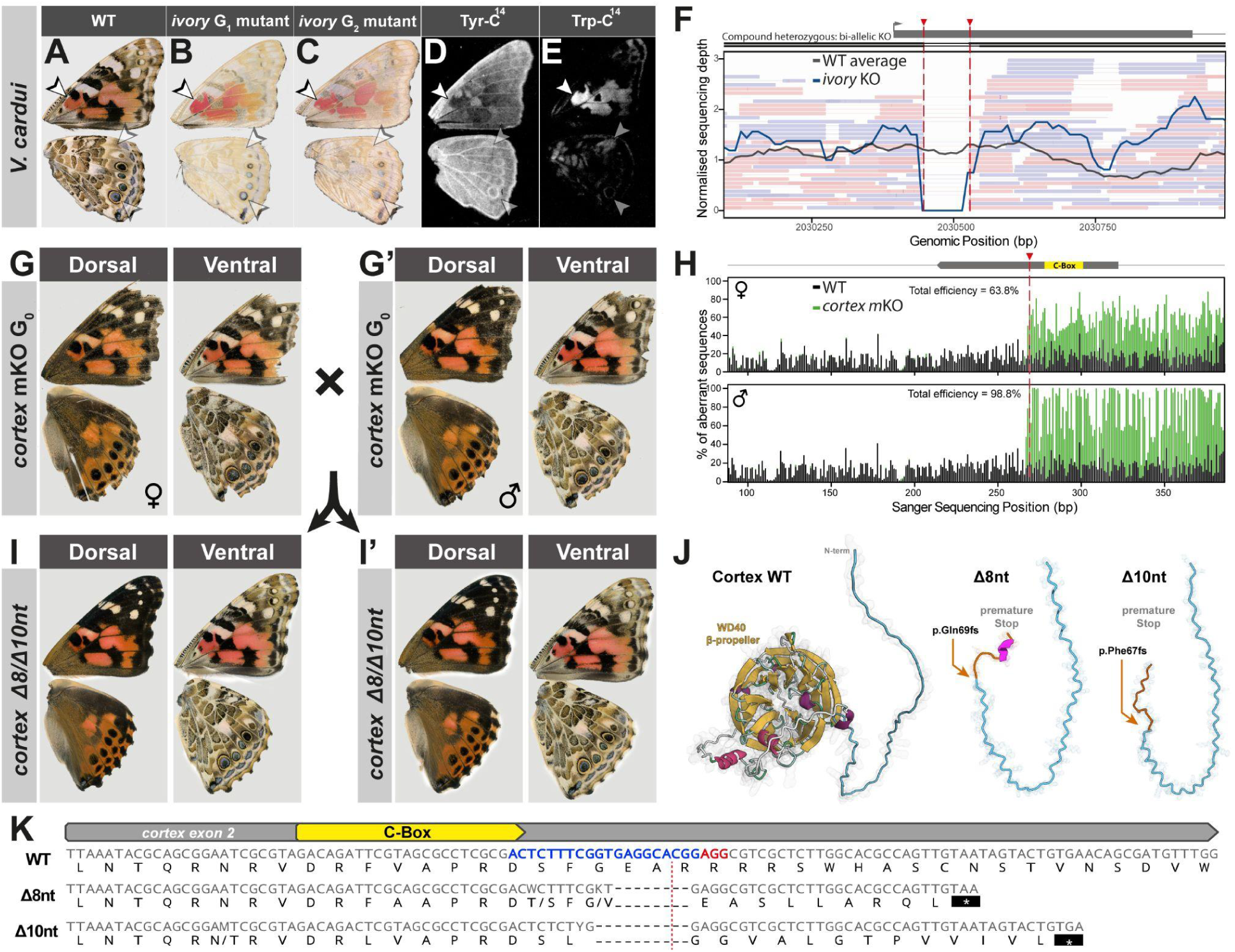
Phenotypes are explained by *ivory* specific mutations. **(A-C)** Loss of melanic scales in *ivory* germline mutants. **(D)** Radiolabeling of Tyrosine highlighting melanin containing scales and **(E)** Tryptophan radiolabeling revealing the distribution of ommochrome-containing patterns. Ommochromes are not affected by *ivory* KOs in *V. cardui* (arrowheads). **(F)** Strong *ivory* phenotype shown in (B) is explained by a compound heterozygous deletion of 82/97 bp at its first exon. **(G-H)** Exon 2 G_0_ *cortex* crispants display no wing phenotypes, but a large fraction of cells carrying mutations at the expected cut site. These crispants were crossed to produce F_1_ offspring **(I-I’)**, again displaying no wing phenotypes. **(J-K)** Genotyped alleles recovered in the F_1_ lead to premature protein truncation due to an 8 nt and 10 nt deletion at *cortex* exon 2.

### Germline mutants of *cortex* rule out a colour patterning function

Previous knock-outs of *cortex* exons generated rare phenotypes in mosaic G_0_ butterflies (6, 9, 10). To circumvent the limitations of mosaic KOs and formally test a function of *cortex,* we generated *V. cardui* null mutants targeting the second exon of *cortex*, an obligatory exon shared between all larval and pupal isoforms in *Heliconius* (**Fig. S1-S2**; Nadeau et al. 2016), and that includes a conserved C-Box motif crucial for APC/C interaction (Livraghi et al. 2021). The resulting *cortex* crispants were indistinguishable from wild-type controls, in spite of high mutation rates observed in the genotyped haemolymph of 7 G_0_ individuals (**Fig S11**). We next crossed two confirmed G_0_ mutants (**Fig. 3G,G’**) and produced F_1_ individuals, again displaying no pigmentation changes on their wings (**Fig. I-I’**). We recovered several compound heterozygous mutants, harbouring two null alleles with premature stop codons, resulting in the early truncation of the *cortex* protein and a complete loss of secondary structure (**Fig. 3J-K and Fig. S12**). Together, these results rule out a function for *cortex* in *V. cardui* wing scale pigmentation.

Importantly, the lack of a *cortex* wing patterning function suggests that *ivory* does not have a local *trans*-regulatory effect on its neighbouring gene, and that the *ivory* promoter is not acting in *cis* to regulate *cortex*. To further examine further potential local *cis*-acting targets, we re-analysed Hi-C and histone ChIP-seq data previously generated in *H. erato* pupal wings (36, 37). Chromatin conformation capture assays detect a strong topologically associated domain (TAD) centred around *ivory* (**Fig. S13**), with no evidence of long-range interactions outside of this domain. Furthermore, histone modification profiling performed in three day old pupal wings reveals that the *ivory* 5’ region coincides with a strong H3K4me3 signal, a hallmark of transcriptionally active promoters (38, 39). Given the absence of other protein-coding genes within the TAD other than *cortex*, and a chromatin state consistent with a promoter rather than enhancer activity, it is likely that *ivory* is acting in *trans*, and not through *cis*-regulation of nearby genes.

### A conserved function for *ivory* in nymphalid butterflies

The sequence conservation of lncRNAs is often limited to 5’ ends of transcripts and degrades rapidly over evolutionary time (40). We identified the *ivory* promoter and first exon as conserved in ditrysian Lepidoptera (**Fig. S14**), which enabled the expression profiling and CRISPR mutagenesis of a homologous region in a further two nymphalid butterflies spanning ∼80 million years of evolution (41). CRISPR/Cas9 targeting of *ivory* in *Agraulis incarnata* and *Danaus plexippus* resulted in crispants displaying scale transformations from melanic to yellow-white states, as well as lighter orange colours (**Fig. 4**).

**Figure 4.**
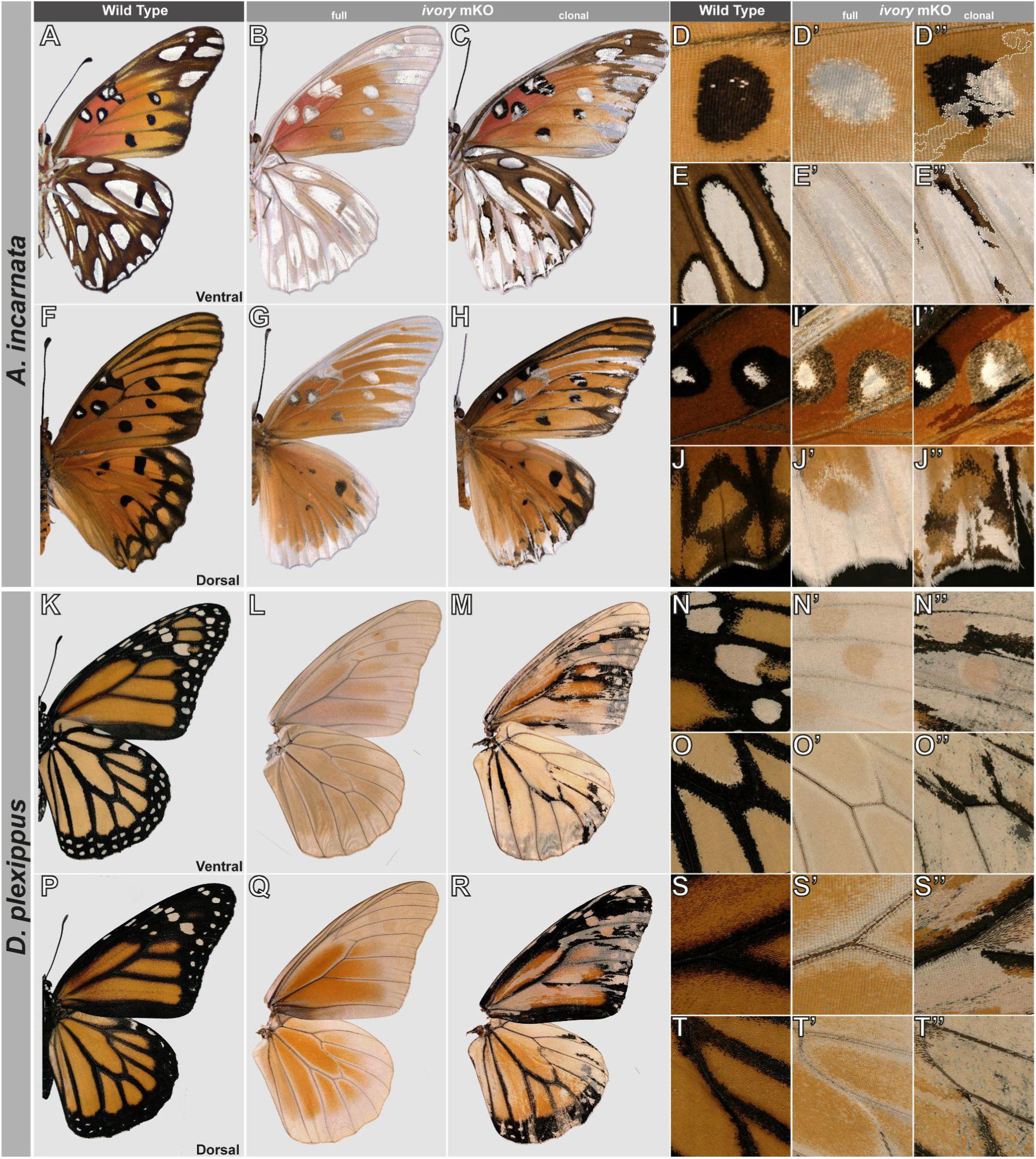
Conserved and pattern-specific effects of *ivory* mKOs on nymphalid butterflies. Wing pattern effects of *ivory* perturbation in representative *A. incarnata* and *D. plexippus* G_0_ crispants. **(A-E’’)** Ventral wing views of *A. incarnata* comparing wild-type to full (non-mosaic) *ivory* crispant and a clonal *ivory* crispant (*i.e.* with fragmented mosaic of WT and mutant clones). Phenotypes include melanic (black/brown)-to-silver transformations and lighter orange (dashed lines). Insets feature magnified views of the Cu_1_-Cu_2_ silver spots in forewings **(D-D”)**, and hindwings **(E-E’’)**. **(F-J’’)** Dorsal views, featuring *ivory* crispant clones with lightened orange and black colouration (**I-I’’**, Discalis II spots), and melanic-to-white conversions (**J-J’**’, marginal M_3_-Cu_2_ region). **(K-O’’)** Ventral wing views of *D. plexippus* comparing wild-type to full and clonal *ivory* crispants. Insets feature the forewing M_1_-M_3_ white spot region **(N-N’’)** and the hindwing discal crossvein **(O-O’’)**, with conversions of both melanic and orange to white states, as well as a residual orange colouration in some of the affected areas. **(P-T’’)** Dorsal views, with insets featuring the forewing Cu_1_-Cu_2_ vein junction **(S-S’’)** and the hindwing M_1_-M_2_ vein junctions **(T-T’’)**.

In *A. incarnata, ivory* mKOs resulted in scale transformations from melanic to reflective white/silver scales (**Fig. 4A-C, Fig. S16**). Closer inspection through scanning electron microscopy revealed that these transformed scales not only underwent a loss of melanin pigmentation, but also a change in the ultrastructure of their top surface (**Fig. S17**), showing an ectopic lamination characteristic of reflective scale types (42). While a subset of dorsal melanic spots only showed a lighter intensity in *ivory*-deficient clones (**Fig. 4G, I-I’’**), others showed complete conversion to white, such as the marginal patterns and venous black markings (**Fig. 4J-J’’**). Orange patterns showed a lighter colouration in crispant clones, likely due to a reduction in melanin content, but consistent with a model where these patterns also incorporate ommochrome pigments that are unaffected in *ivory* mKOs (**Fig. S16**).

In *D. plexippus ivory* crispants, melanic scales again turned white, and orange scales also appeared lighter, an effect especially pronounced on ventral wing surfaces (**Fig. 4K-M**). Interestingly, distal white spots on the ventral forewings changed from white to orange, while more distal orange areas showed the opposite effect, turning from orange to white, indicating that there are local differences in *ivory* function (**Fig. 4N-N’’**). These effects were consistent on both wing surfaces in *D. plexippus* (**Fig. 4P-R**), with pronounced melanic-to-white around the vein margins in both forewings (**Fig. 4S-S’’**) and hindwings (**Fig. 4T-T’’**).

## Discussion

### Cortex – a case of mistaken identity?

Colour variation loci in peppered moths and *Heliconius* butterflies include the protein coding gene *cortex*, which was originally proposed as a causal gene within the corresponding genetic intervals based on its differential expression profile (1, 4). Several follow-up studies used CRISPR to mutagenize a coding exon of *cortex*, and generated crispants with discoloured clones in *Heliconius*, *Danaus*, *Junonia*, and *Kallima* butterflies (6, 9, 10), further anchoring the incorrect impression that *cortex* itself is necessary for colour specification during development. In parallel with findings in *Junonia coenia* (*in submission*), we show that *ivory*, a lncRNA overlapping with *cortex,* is in fact the causal gene.

This misidentification was caused by two main issues. First, *ivory* had escaped detection as it was missing from former genome annotations. This is likely the consequence of the erroneous exclusion of non-coding transcripts by genome annotation pipelines, and is reflective of a general bias towards protein coding genes in our understanding of development and evolution. Second, CRISPR repair events occasionally create deletion alleles that are much larger than intended (43–48). We also recovered large deletions in the 10kb-30kb range in a batch of G_1_ siblings following CRISPR targeting (**Supporting Text and Fig. S18**). Assuming that CRISPR-induced structural variants occur at low frequency, we suggest that sporadic, large deletions induced by *cortex* mutagenesis fortuitously reached *ivory*, explaining the low penetrance of wing phenotypes in these previous experiments. It thus appears unlikely that *cortex* modulates pigment variation in other butterflies and moths. Instead, the *ivory* lncRNA, which is partly nested within the *cortex* gene, is a proven regulator of colour scale specification and the likely causal driver of pattern variation in natural populations.

### A lncRNA repeatedly drives adaptive phenotypic variation

There has been considerable interest in the *cortex/ivory* region, as it was independently mapped as the master switch locus behind striking examples of adaptive variation, namely industrial melanism in peppered moths (1, 2), Müllerian mimicry in *Heliconius* (3–7), Batesian mimicry in *P. clytia* (8), seasonal polyphenism in *Junonia* (10), and leaf-mimicking crypsis in oakleaf butterflies (9). Adaptive variation between morphs consists of shifts in the intensity and spatial distribution of melanin-containing scales in all of these cases. In our study, we showed that *ivory* expression consistently prefigured the distribution of melanic scales in nymphalid butterflies, and that in fact *ivory* and not *cortex* is required for their specification.

This stimulates a reframing of previous population genetics studies. We now infer that regulatory alleles of *ivory* drive the repeated evolution of pattern variation involving a shift between black and white/yellow. Indeed, *H. melpomene* and *H. erato* co-mimics geographically vary in the display of their hindwing yellow-band and marginal white-spots across their range. These polymorphisms map to modular non-coding regions of the *cortex-ivory* locus (**Fig. 1B**), that independently control melanism over the ventral and dorsal hindwing yellow bars, and the white hindwing marginal regions (4, 6, 13, 25–27). As *ivory* controls melanic pattern distribution in *Heliconius* (**Fig. 2A,B**), these population genomic signals likely point at *cis*-regulatory elements that switch *ivory* expression on or off at these colour patterns. Likewise, in *Heliconius numata* and *Kallima inachus*, adaptive colour morphs have been linked to inversion polymorphisms, allowing complex regulatory haplotypes to accumulate in the absence of recombination (7, 9, 49). This mechanism likely underlies the fine-tuning of *ivory* expression, and while a role of other genes within these inversions is not excluded (*ie.* a supergene), *ivory* likely explains a large fraction of the colour pattern variations that are involved in Müllerian and leaf mimicry in these species.

### The *ivory* lncRNA promotes melanic scale identities

The expression of *ivory* prefigures the position of melanic patterns at a stage where known master selector genes define scale identity (50, 51), and perturbation of *ivory* resulted in the transformation of melanic scales into white or light-coloured identities. Melanic scales contain either NBAD-melanin or DOPA-melanin pigments; two end-products of the insect melanin synthesis pathway following the metabolism of Tyrosine into dopamine pigment precursors (52, 53). Lighter coloured scales may contain the dopamine byproduct NADA-sclerotin, a clear molecule that crosslinks cuticular proteins and participates in scale hardening (54, 55). Our working model thus posits that *ivory* activates melanic colour outputs by indirectly promoting dopamine-melanin synthesis or by antagonising the production of NADA-sclerotin, explaining the predominance of white scales in *ivory* knock-outs. Modulating the ratio of dopamine-melanin and NADA-sclerotin may also explain the ultrastructural increase in the upper lamina of *ivory-*deficient scales (**Fig. S17**), since similar effects on scale upper laminae occur in *Bicyclus* mutants with reduced dopamine-melanin (52). Of note, some areas in *A. incarnata* and *V. cardui* crispants showed partial phenotypes, with orange patterns showing a lighter colouration in mutant clones. This effect is likely explained by a mix of pigment in these scales, supported by radiolabeling experiments (**Figs. S7-S9, S16**). These data suggest that while nymphalid colour patterns can consist of discrete melanic and ommochrome identities, some scales can also contain mixtures of both pigment types.

Finally, *ivory* crispant clones also showed a light-yellow pigmentation in *H. erato* and *H. charithonia* (**Figs. S8-S9**), which is attributed to 3-hydroxykynurenine (3-OHK), a yellow Trp-derived pigment that is incorporated into yellow *Heliconius* scales from circulating hemolymph during late pupal development, and not into black scales (56–58). We thus conclude that *ivory* acts to prevent 3-OHK incorporation into melanic scales (**Fig. S9D-F**). Taken together, these results suggest that *ivory* acts as an upstream regulatory factor that can orchestrate several aspects of scale pigmentation, not only by promoting dark melanin metabolism, but also by blocking 3-OHK incorporation.

The precise molecular mechanism by which *ivory* acts is not yet clear. lncRNAs have been described as participating in a wide range of biological functions, from *cis-*effects on transcription initiation and elongation, to the regulation of chromatin conformation, to *trans*-effects on transcription and translation of other genes (19, 23, 59). Here, we have shown *ivory* is not acting in *cis* to regulate the adjacent *cortex* gene, as deletion of the protein coding region of *cortex* does not affect pattern, and as chromatin conformation and histone-modification at the *ivory* promoter is inconsistent with a distal enhancer role function (**Fig. S13**). Therefore, we infer that *ivory* likely acts in *trans* to regulate other genes or processes.

## Conclusion

Only a few lncRNA genes have been linked to adaptive variation in natural populations, including a locus producing phased siRNAs that modulate monkeyflower colouration, and a nuclear RNA involved in thermal adaptation in fruit flies (60, 61). In this study, we showed that the *ivory* lncRNA is spatially regulated during development and is required for melanic scale identity. Remarkably, this role is conserved by nymphalid butterflies across 80MY of divergence (41), and includes cases where allelic variation at the *ivory* locus itself drives phenotypic adaptations, such as the yellow band vs. melanic pattern switches involved in *Heliconius* mimicry. Future characterization of *ivory* alleles in polymorphic populations of butterflies, as well as in geometrid moths that underwent episodes of industrial melanism (2, 62), promise to be an exciting avenue of research on the molecular basis of adaptation. More generally, our findings further establish that lncRNAs are not only important regulators of development, but also pivotal drivers of evolutionary change at the genetic level.

## Methods

### Butterflies

*H. erato demophoon* and *H. melpomene rosina* (origin: Panama), *and H. charithonia charithonia* (origin: Puerto Rico) were reared in greenhouse environments using primarily *Passiflora biflora* as a host plant, as well as *Passiflora menispermifolia* and *Passiflora vitifolia* for *H. melpomene* (6). *A. incarnata* were reared at 25°C or 28°C on *P. biflora*, *Passiflora caerulea*, *Passiflora incarnata* or an artificial diet (passionvine butterfly diet, Monarch Watch Boutique, supplemented with 30 g/L of water of fine ground *P. biflora* leaf powder). *A. incarnata* adults were kept in a greenhouse cage with nectaring sources (Gatorade 50% in feeding cups, *Lantana camara, Buddleja davidii*) and supplemented with UV-light (Repti-Glo 10.0 Compact Fluorescent Desert Terrarium Lamp bulbs, Exo Terra). *V. cardui* (stock origin : Carolina Biological Supplies) were reared on artificial diet with oviposition on *Alcea rosea* or *Malva sylvestris* (63, 64). *D. plexippus* were reared on *Asclepias curassavica* in greenhouse conditions.

### *De novo* transcriptome assemblies

Raw RNA-seq data reads were obtained from Bioproject PRJNA552081 (Hanly et al. 2019) and used to generate *de novo* transcriptome assemblies for *H. erato demophoon*, *H. erato hydara*, *H. melpomene melpomene* and *H. melpomene rosina* (Table 1). Transcriptome assemblies were generated using Trinity v2.10.0 with a minimum length of 200 bp and the default K-mer of 25 (65). Residual adapters present in the assembly were trimmed using FCS adaptor (66) and contigs less than 200 bp in length were removed.

### CRISPR mosaic knock-outs (mKOs)

Butterfly embryo micro-injections followed published procedures (6, 63, 67) using 1:1 or 2:1 mass ratios of Cas9-2xNLS (PNABio or QB3 Macrolabs) and synthetic sgRNAs (Synthego, listed in **Table S3**). Injections were completed before blastoderm formation around 4 h (**Table S1**), most of them within 2 h AEL, thus preceding the first mitotic cleavage (68–70). Survival and phenotypic penetrance were assessed in neonates, pupae, and adults, and no morphological phenotypes were visible in immatures in this study.

### *In-situ* hybridizations

Chromogenic RNA *in-situ* hybridization (ISH) followed previously described procedures (71, 72) with the following modifications. Antisense riboprobes targeting the conserved *ivory* first exon of each species were transcribed from PCR templates (ref), that were amplified from wing cDNA and gDNA (**Table S4**). Developing wings were dissected at 20-35% pupal development, incubated in fixative for 30-40min (1X PBS, 2 mM egtazic acid, 9.25% formaldehyde), and stored in MeOH at −20°C. On the day of the hybridization procedure, wings were rehydrated progressively in PBT (1X PBS, 0.1% Tween20), and digested with 2.5 µg/mL Proteinase K for 5 min on ice before post-fixation and hybridization with 40 ng/µL of DIG-labeled riboprobe at 60-63°C. Wings were stained at room temperature for 4-6 h in BM Purple (Roche Applied Science).

### Data, Materials, and Software Availability

Whole-genome sequencing data are available in the Sequence Read Archive (www.ncbi.nlm.nih.gov/sra) under BioProject accession numbers PRJNA1031670. Trinity assemblies have been deposited on the open science network repository under URL: https://osf.io/q3sy7/?view_only=ef779e69225443b3a5aae7ddfe1f292a.

## Acknowledgements

We thank T. Kirby and R. Canalichio for help with butterfly husbandry; S. Van Belleghem for bioinformatics assistance; the GWU HPC team for providing computing infrastructure; the GW Nanofabrication and Imaging Center (GWNIC) for microscopy resources. This work was supported by the following awards: NSF IOS-2110534 to AM; NSF EPSCoR OIA-1736026, PRSTRT ARG 2020-00138, and funding from the FIPI at the UPR-RP to RP.

## Author contributions

Conceptualization, LL, JJH, CDJ and AM; Methodology, LL and JJH; Investigation, LL, JJH, EE, CJW, LSL, AM-V, KK, AC, ESMH, RDR, and AM; Writing, LL, JJH and AM; Review & Editing, LL, JJH, AM; Funding Acquisition and Resources, JJH, RP, CDJ, and AM.

## Competing interest statement

The authors declare no conflict of interest.

## Classification

Biological Sciences: Evolution

## Supplementary Information

### Supplementary Methods

#### RNA-Seq analysis

Previously published *H. melpomene* and *H. erato* RNA-seq reads (NCBI SRA: PRJNA552081) (28) were realigned to their corresponding genomes (Hmel v2.5 and Hera v1.0), downloaded from LepBase (73). Reads were aligned using the splice-aware STAR aligner v2.7.10a (74), taking into account strandedness information. The resulting BAM files were then visualised in IGV, and strand specific junctions were identified using the inbuilt junction track function. Spurious splicing events were filtered to exclude any events that occurred with less than 10 supported reads. The combined evidence from the aligned STAR results and the Trinity assemblies were used to annotate the *ivory* transcript in the *Heliconius* genomes. Normalised counts and differential expression for RNA-seq data were calculated using the R package DESeq2 (75) comparing samples by developmental time, with significance reported as adjusted *p*-values.

#### ATAC-Seq analysis

Previously published *H. melpomene* and *H. erato* ATAC-seq reads (SRA: PRJEB43672) (6) were realigned to their corresponding genomes (Hmel v2.5 and Hera v1.0) using bowtie2 v.2.5.0. Normalised counts and differential accessibility were calculated using the R package DESeq2 (75) comparing samples by developmental time, with significance reported as adjusted *p*-values. ATAC-Seq tracks were plotted by subtracting head derived reads from a normalised read depth average calculated from hindwing posterior compartments of all morphs at 36 h after pupa formation.

#### 3’ RACE for V. cardui ivory

3’ RACE (Rapid Amplification of cDNA Ends) was performed using RNA from pupal forewing and hindwing tissue at 15%, 30% and 45% of development, using the Superscript IV First-Strand Synthesis System kit (Invitrogen) for cDNA generation, and following published recommendations (76). Briefly, a gene specific primer anchored to *ivory* exon 1 was used for first round amplification (**Table S5**). Secondary rounds of amplification were then performed by using a further nested primer on exon 1 using the first PCR product as a template. Isoform products were extracted from a 1% agarose gel using the Zymoclean Gel DNA Recovery Kit (Zymo Research) before Sanger sequencing.

#### Whole-genome sequencing and genotyping of G_1_ and G_2_ *V. cardui ivory* mutants

Syncytial embryos collected from *V. cardui* were injected between 45 min and 2 h AEL, using a combination of two sgRNAs targeting either the first or second exon of *ivory* (**Tables S1 and S3**). Adult crispants displaying large mutant clones in the G_0_ were pooled in cages of 4-15 butterflies for random in-crossing. G_1_ individuals were randomly in-crossed again in cages of 4-15 butterflies to generate G_2_ individuals. DNA from *V. cardui* butterflies displaying strong *ivory* phenotypes was then extracted from adult thoraces. Sequencing libraries (PE 150) were generated and sequenced in an Illumina NovaSeq6000 S4 Flow cell at the Institute for Genome Sciences (University of Maryland -Baltimore). Read alignments to the *V. cardui* reference genome (77) were generated using BWA-MEM (78). Local sequencing coverage was determined using Mosdepth (79). To normalise the data, coverage values for each window were divided by the average depth for the entire scaffold. Wild-type coverage was calculated by averaging the median normalised counts across 4 control *V. cardui* individuals, and used to compare to the G_1_ and G_2_ normalised counts, to account for possible population specific structural variation (**Fig. 2H and Figs. S11 and S18**).

#### Autoradiographs of wing pigment precursors

Pupae were injected with radiolabeled tryptophan or tyrosine during late pupal development, at a stage when ommochrome pigmentation just becomes observable in the eyes. Pupae were injected in the abdomen with a Hamilton syringe with approximately 0.1 μCi of L-[methyl-^14^C]-tryptophan or L-[^14^C(U)]-tyrosine (New England Nuclear Corporation, MA) pre-diluted in a sterile saline solution (130 mM NaCl, 5 mM KCl, 1 mM CaCl_2_, pH = 6.9). Adult butterflies were allowed to eclose, and frozen within 12 hours of emergence. Scales from dorsal and ventral wing surfaces, respectively, were removed using adhesive polypropylene packing tape, which was then covered on the adhesive side using polyethylene food wrap to seal the scale print. Mounted scale prints were exposed to HyBlot CL autoradiography film (Denville Scientific, Metuchen, NJ), with the polyethylene wrap side against the film, in a −80°C freezer for 9 d (*V. cardui*) or 12 d (*H. erato*). Autoradiograms were developed on an automated film processor and digitally scanned.

#### PCR genotyping of G_0_ and G_1_ *V. cardui cortex* mutants

CRISPR injections used a single sgRNA targeting the obligatory second exon of the *cortex* transcript, containing the conserved C-box motif (**Table S3**). Resulting G_0_ individuals were genotyped at the larval or pupal stage using a non-invasive method. In short, 1 µL of haemolymph was sampled and used for PCR amplification of the *cortex exon 2* (**Table S5**) using the diluted protocol of the Phire Tissue Direct PCR Master Mix (Thermo Fisher Scientific), before PCR purification using the DNA Clean & Concentrator Kit (Zymo Research), and Sanger Sequencing. Chromatograms were used for trace decomposition using *TIDE* (80), in order to identify butterflies with high frequencies of indel alleles at the sgRNA cutting site. Confirmed G_0_ butterflies were crossed in single pairs to generate a F_1_ generation. Siblings from a single cross were PCR amplified and sequenced, and mutant alleles at heterozygous and compound heterozygous states were extracted using *TIDE* and *Poly peak parser* (80, 81). The structure of WT and inferred frameshifts protein sequences of *V. cardui* Cortex was simulated using AlphaFold Colab and visualised in Mol* Viewer (82, 83).

#### Chromatin profiling

Raw Hi-C reads for *Heliconius erato lativitta* whole pupal wings were obtained from NCBI BioSample SAMN10587321. The downloaded Hi-C sequencing reads were aligned to the *Heliconius_erato_lativitta_v1* genome assembly using bwa-mem (84). The HiCExplorer v3.7.3 suite was employed to identify restriction sites specific to the DpnII enzyme (85). A Hi-C matrix was built based on the identified restriction sites using merged counts from two independent replicates for each wing tissue (Forewing and Hindwing). Datasets for forewings and hindwings were normalised to the smallest given read number. The matrix underwent correction using a matrix balancing algorithm (86). HiCExplorer employs a TAD-separation score to assess the extent of separation between each side of the sliding bins within the Hi-C matrix, with bins set at 10 kb in this instance. Frequency contacts were visualised using hicPlotMatrix and hicPlotTADs commands. Previously published chromatin accessibility profiles, including ATAC-seq and histone modification ChIP-seq datasets, were downloaded from NCBI GEO dataset GSE111023, GSE109889 & GSE105080 (36).

#### Scanning Electron Microscopy (SEM)

Sample preparation and imaging was conducted according to a published procedure (42). Briefly, cut wing samples were mounted onto a stub using a double-sided carbon tape. Images of mounted stub were taken using a Keyence VHX-5000 microscope before sputter-coating with 8 nm of gold using a Cressington 208HR Sputter Coater. SEM images were acquired using an acceleration voltage of 2.00 kV, a beam current of 25 pA and 10 μs dwell time.

### Supplementary Figures

**Figure S1.**
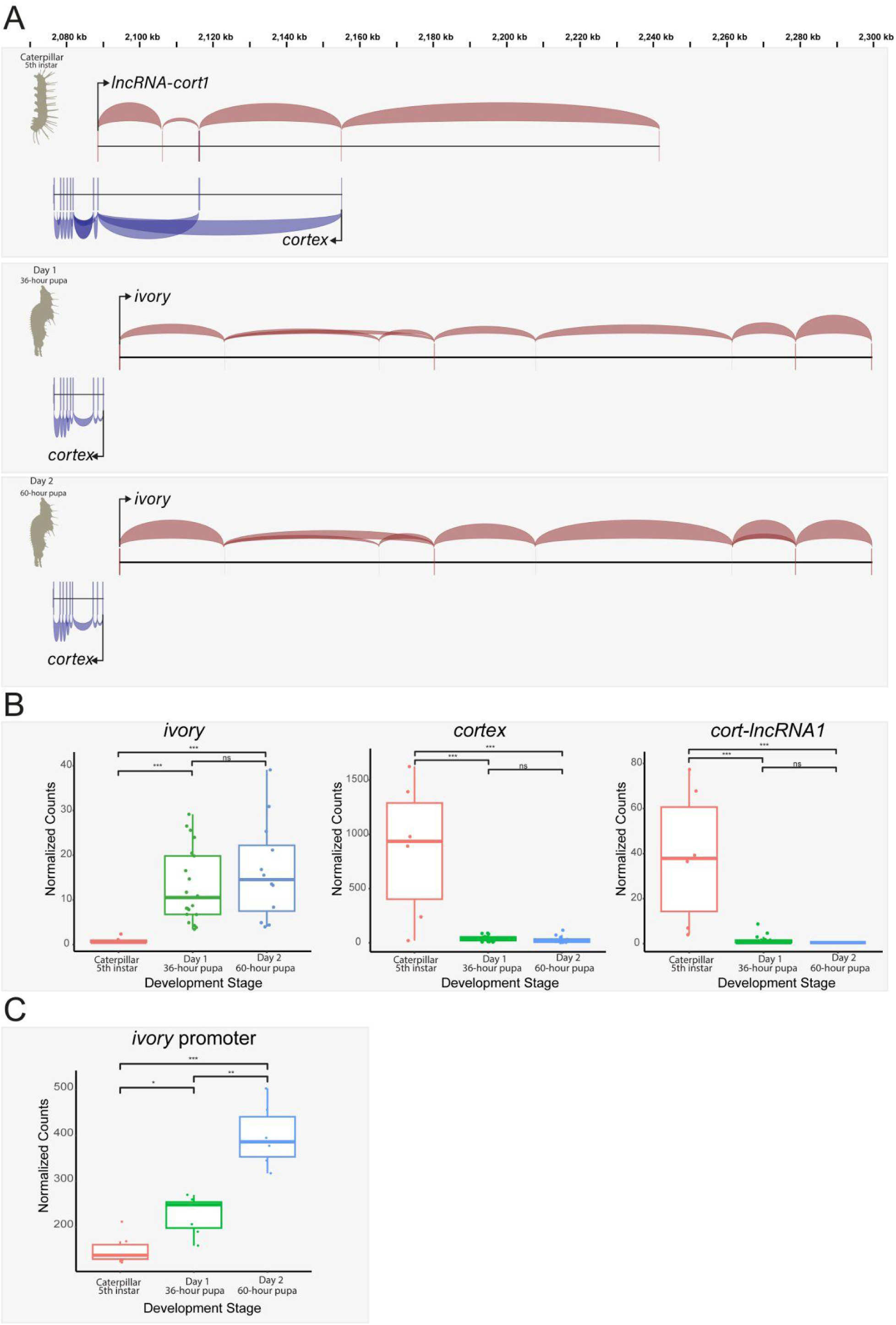
Absence of *ivory* expression in 5th instar wings is followed by an increase during pupal wing development in *H. erato*, concomitant with an increase in promoter availability. **(A)** Annotations and splicing events recovered from the stage specific RNA-seq dataset are shown for lncRNAs *lncRNA-cort1* and *ivory* (red) and *cortex* (blue), showing *ivory* is a pupal wing specific transcript. **(B)** Expression profiling of transcripts of the *cortex* locus in *H. erato*, based on a reanalysis of published wing RNA-seq (Hanly et al. 2019; Livraghi et al. 2021) indicating an increase in *ivory* expression over time, concurrent with a decrease in *cortex* and *lncRNA-cort1* expression. Expression levels are plotted as DESeq2 normalised counts. Pairwise Wald tests adjusted for multiple test correction each assess differential expression between developmental stages. ns : non-significant; *** : *p* < 0.001. **C.** Chromatin accessibility at the *ivory* promoter calculated from published wing ATAC-Seq data at the corresponding stage from panel C (Livraghi et al. 2021; van Belleghem et al 2023), showing an increase in promoter availability accompanies an increase in *ivory* expression. Promoter accessibility is plotted as DESeq2 normalised counts. Pairwise Wald tests adjusted for multiple test correction each assess differential expression between developmental stages. ns : non-significant; * : *p* < 0.05; ** : *p* < 0.01; *** : *p* < 0.001.

**Figure S2.**
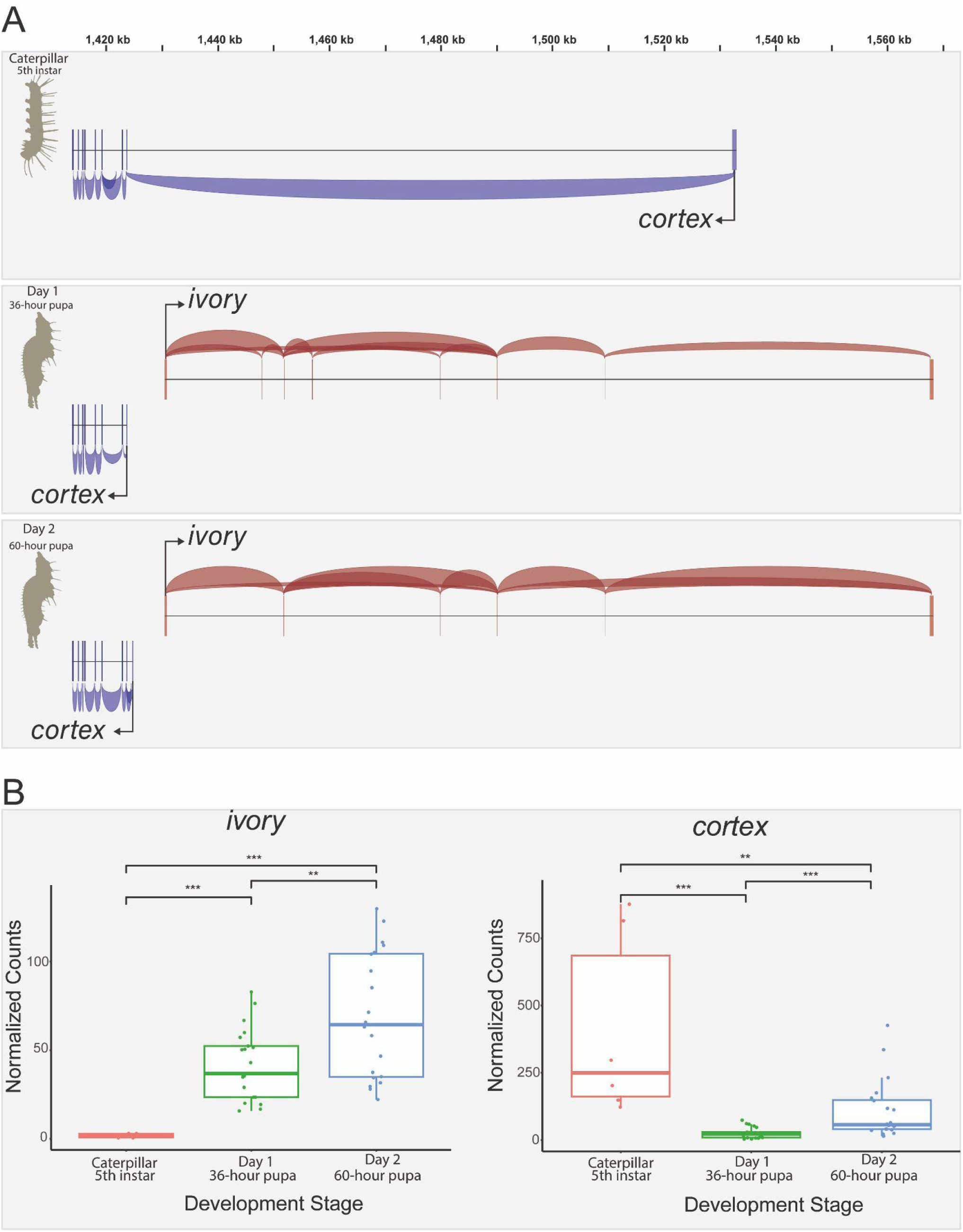
Absence of *ivory* expression in 5th instar wings is followed by an increase in expression during pupal wing development in *H. melpomene*. **(A)** Annotations and splicing events recovered from the stage specific RNA-seq dataset are shown for *ivory* (red) and *cortex* (blue), showing *ivory* is a pupal wing specific transcript. **(B)** Expression profiling of transcripts of the *cortex* locus in *H. melpomene*, based on a reanalysis of published wing RNA-seq (Hanly et al. 2019; Livraghi et al. 2021) indicating an increase in *ivory* expression over time, concurrent with a decrease in *cortex* expression. Expression levels are plotted as DESeq2 normalised counts. Pairwise Wald tests adjusted for multiple test correction each assess differential expression between developmental stages. ** : *p* < 0.01; *** : *p* < 0.001.

**Figure S3.**
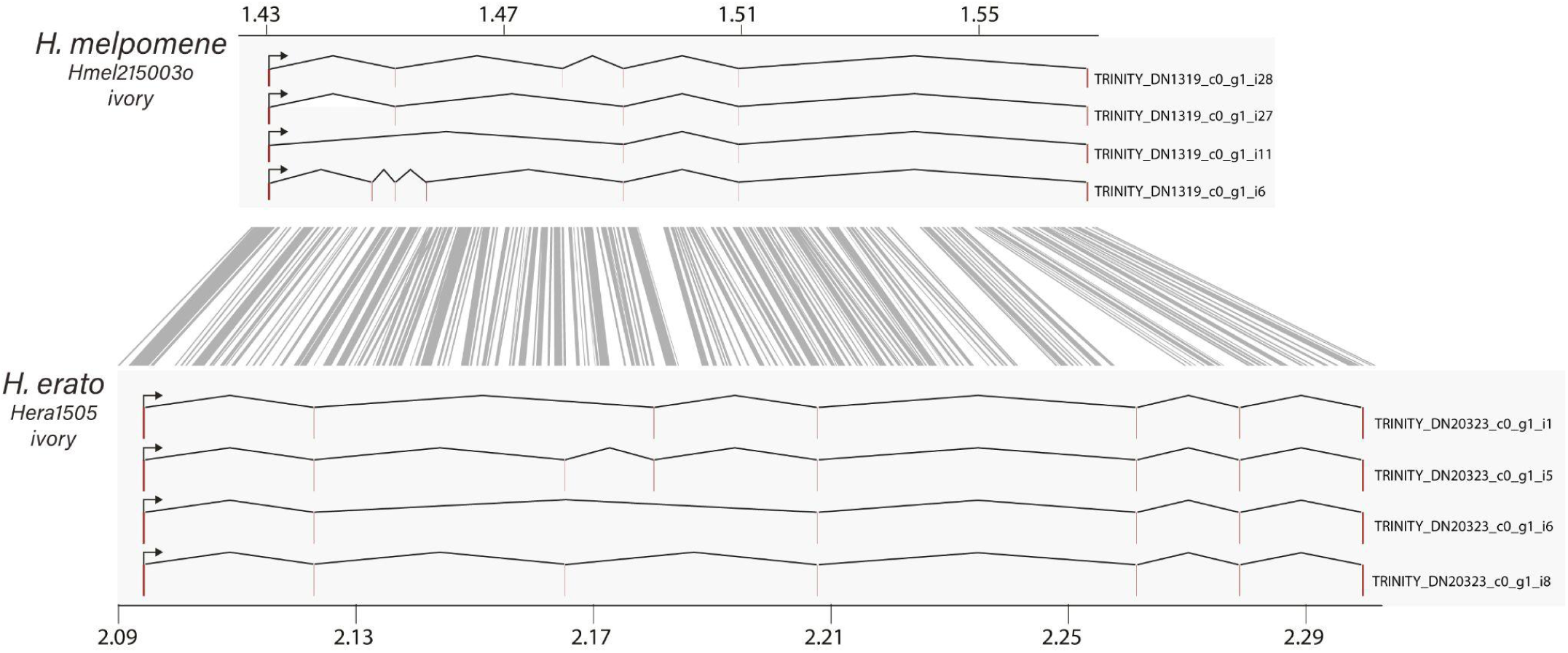
*De novo* transcriptome assemblies identify four *ivory* lncRNA isoforms in *Heliconius*. Assembled transcripts for both *H. melpomene* and *H. erato ivory* from Trinity assemblies, and their FASTA identifiers are shown at their corresponding genomic positions. Full sequences for each of the transcripts can be found in the supporting text. Trinity assemblies are accessible on the Open Science Framework repository under DOI:10.17605/OSF.IO/Q3SY7 (87).

**Figure S4.**
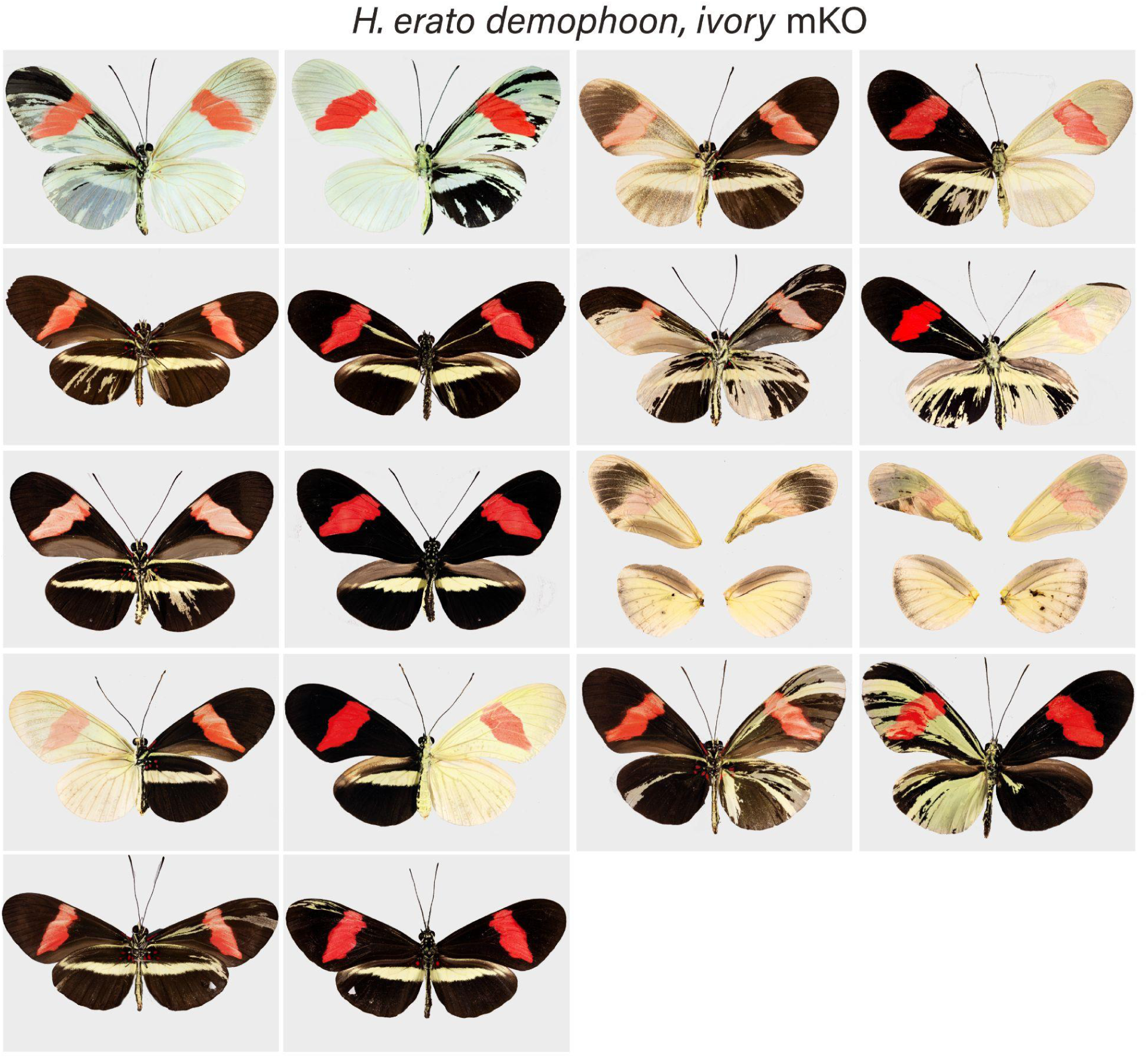
Representative *H. erato demophoon ivory* crispant phenotypes. Ventral (left image) juxtaposed to dorsal (right image) sides for each individual.

**Figure S5.**
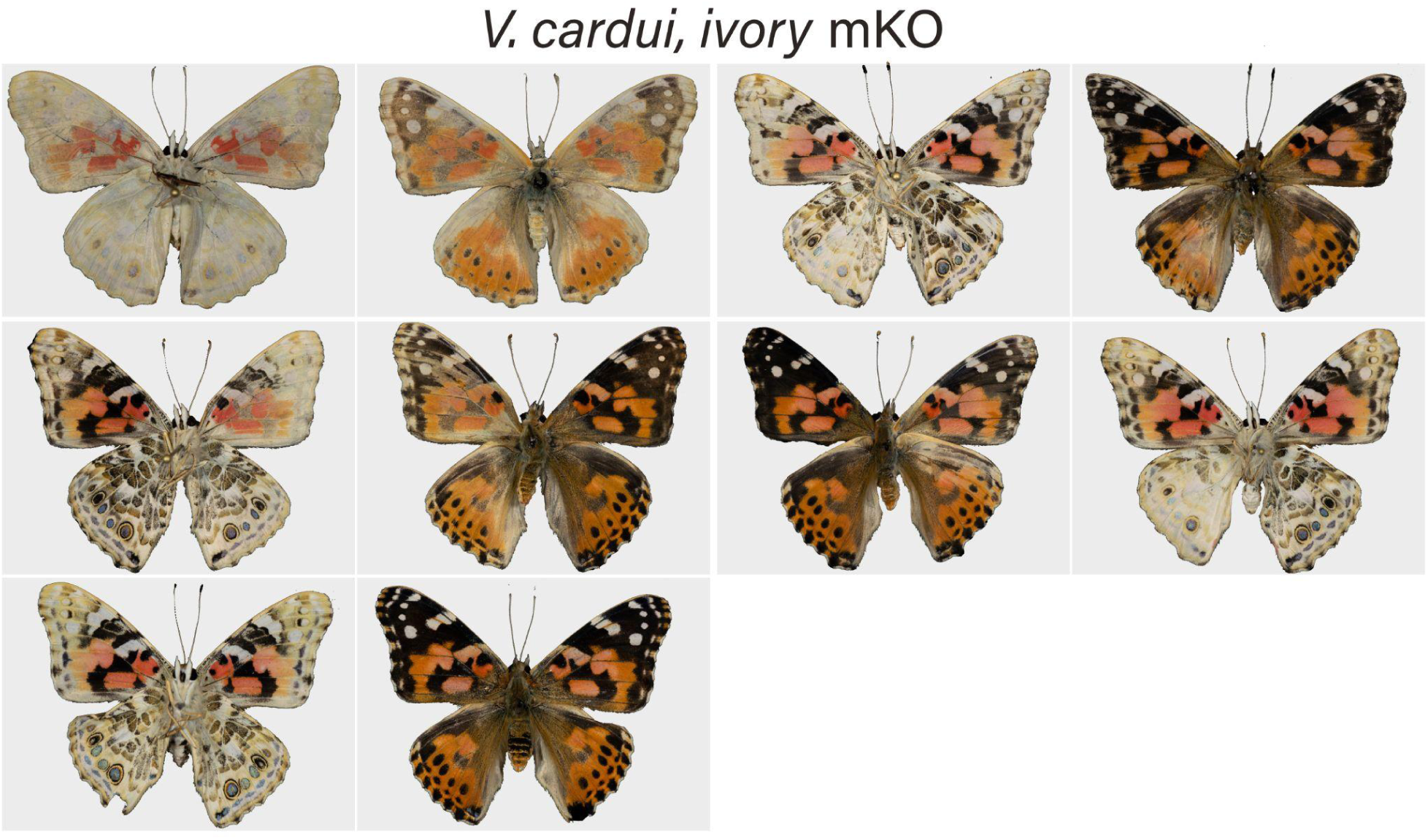
Representative *V.cardui ivory* crispant phenotypes. Ventral (left image) juxtaposed to dorsal (right image) sides for each individual.

**Figure S6.**
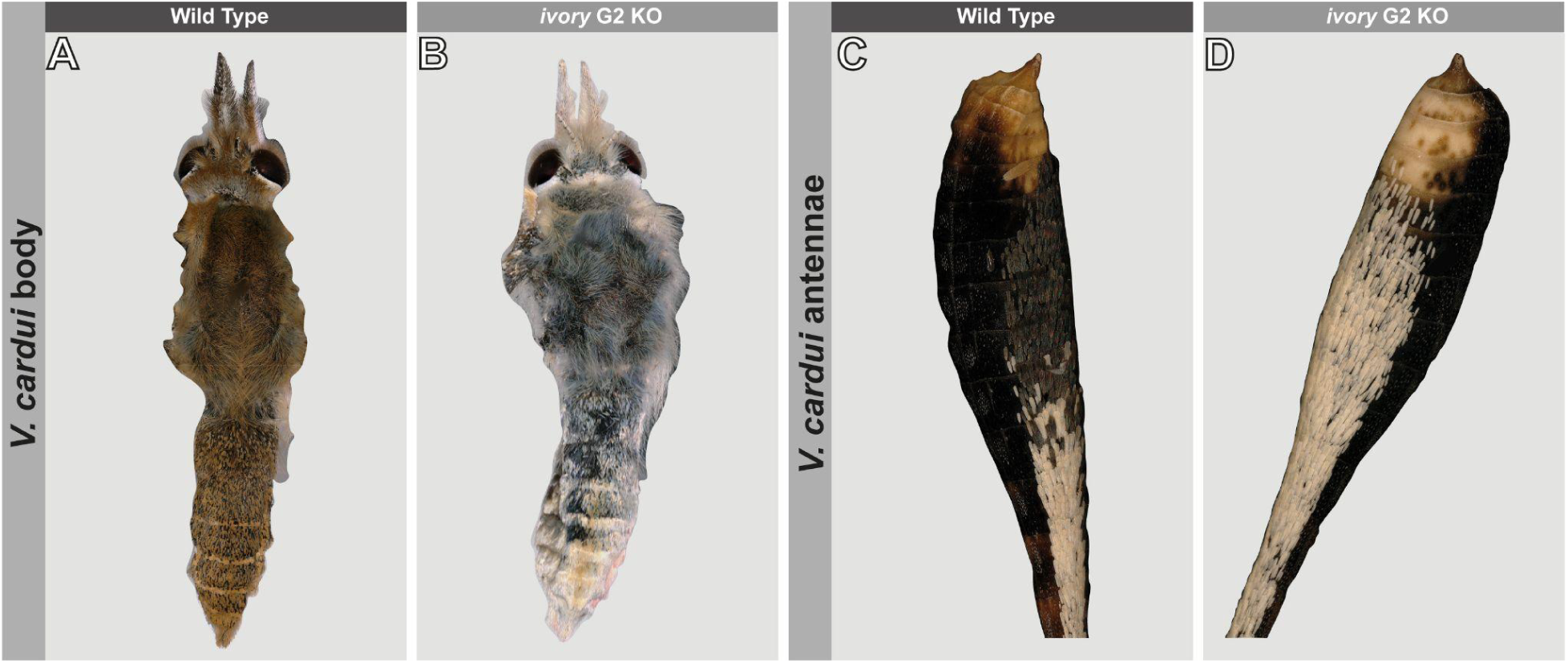
Melanic scale loss in bodies and antennae of *ivory* mutants in *V. cardui*. **(A)** Wild-type adult *V. cardui* showing brown scales on the body and thorax. **(B)** A G_2_ *ivory* mutant displaying a loss of melanic scales on the body. **(C)** Wild-type black scales on the antennal tip and **(D)** transformation to white scales in *ivory* G_2_ mutants.

**Figure S7.**
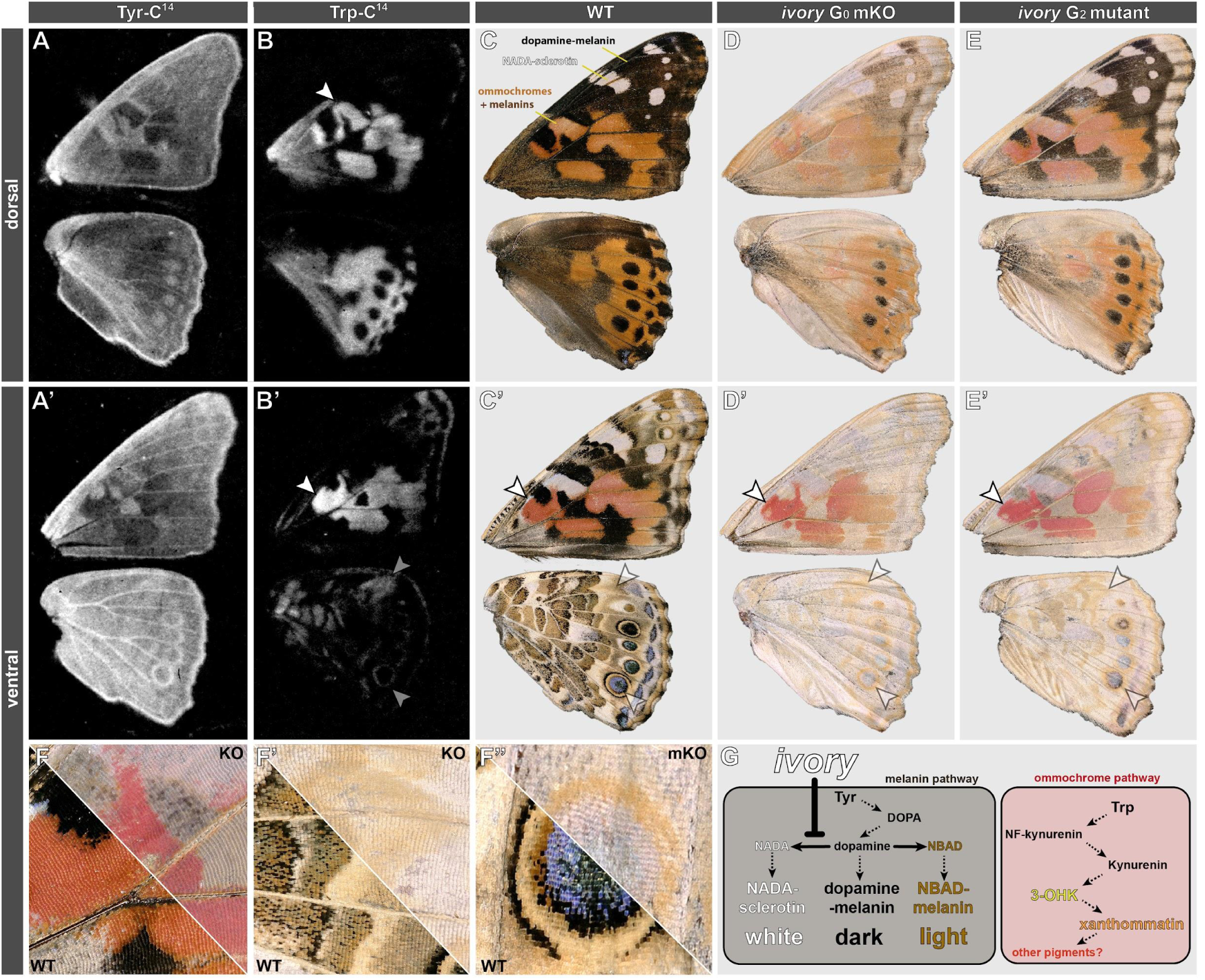
Radiolabelling experiments suggest *ivory* KOs affect melanin but not ommochrome classes of pigments in *V. cardui*. (**A-A’**) Autoradiographs of C^14^-labelled Tyrosine incorporation in the dorsal and ventral wings of V. cardui, highlighting pervasive Tyrosine metabolism across the scales of both surfaces, with increased intensity in presumptive dopamine-melanin (dark) patterns. (**B**) Autoradiographs of C^14^-labelled Tryptophan incorporation, revealing that ommochrome pigment derivatives are strictly limited to orange patterns on the dorsal side. (**B’**) On the ventral side, ommochromes show a strong signal in the orange-pink areas of the ventral forewing (white arrowhead), as well as localised deposition in patterns with beige-yellow tints in the adult (grey arrowheads), including yellow eyespot rings. (**C-E’)** Effects of *ivory* perturbation on *V. cardui* colouration, as shown in WT (C-C’), a G_0_ *ivory* crispant (D-D’), and a G_2_ individual with bi-allelic *ivory* mutations (**Fig. S11D**). (**F-F”**) Magnified views of *ivory*-deficient patterns (as indicated by arrowheads in B-E’). Depigmentation effects are consistent with a pervasive loss of melanin pigments, while putative ommochrome pigments appear to remain in mutants. (**G**) Working model for *ivory*’s role in *V. cardui* pigment modulation, with a conserved inhibitory function on the production of sclerotin (clear pigment). Ommochrome pigments are seemingly independent of *ivory* in this butterfly, as they did not expand or disappear in mutants.

**Figure S8.**
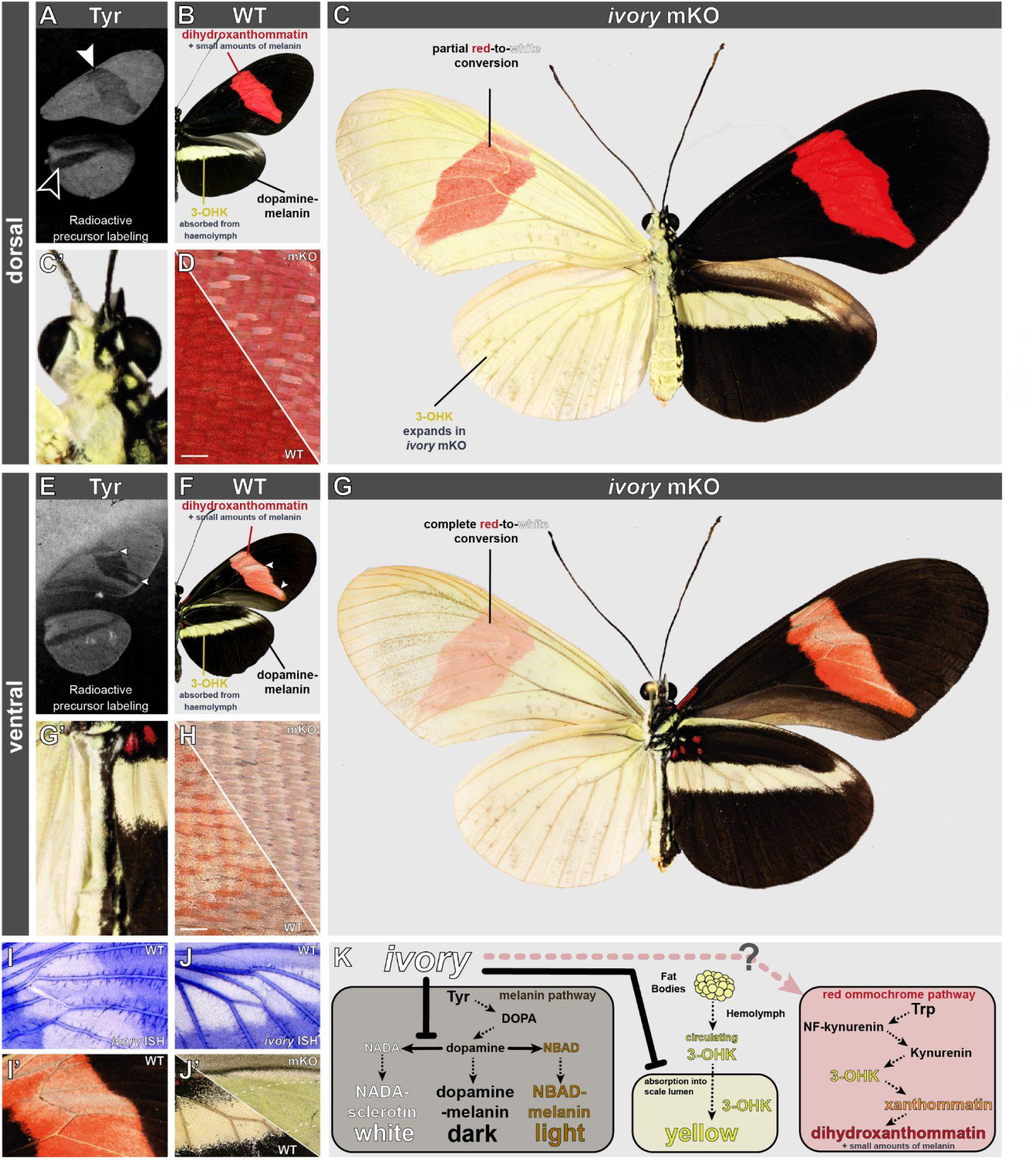
Mosaic knock-outs of *ivory* suggest a regulatory role in *H. erato*. **(A-B)** Dorsal wing autoradiographs marking the incorporation of isotope-labelled Tyrosine in presumptive melanic regions of *H. erato*. 3-OHK, the yellow ommochrome pigment in this species, is known to be incorporated from the hemolymph, while dihydroxanthommatin is the likely red ommochrome pigment (58, 88). **(C-C’)** *H. erato demophoon* G_0_ crispant for *ivory* (shown here in mirrored orientation), with a unilateral colouration phenotype on one side (C’ : magnified view of the dorsal head and thorax). **(D)** Magnified view of red scale phenotypes on the dorsal forewing red bar (WT control vs. *ivory* mKO), with a partial conversion of red-to-white. **(E-G)** Same as panels A-C but with ventral views. Arrowheads : streaks of low Tyr-incorporation in the ventral side of the red forewing band, corresponding to areas characterised by white scales (devoid of ommochrome, but likely incorporating traces of Tyr to make NADA-sclerotin). G’ : magnified view of the ventral abdomen. **(H)** Depigmentation of the ventral forewing red bar. **(I-I’)** Magnified view of a forewing pupal ISH staining of *ivory* lncRNA expression, denoting the presumptive melanic patterns around the red band (adult WT ventral view shown in I’). (**J-J’**) Magnified view, *ivory* lncRNA expression in the hindwing, demarking the presumptive melanic patterns around the yellow stripe. **(K)** Working model for *ivory* functions, with a block of both 3-OHK incorporation and NADA-synthesis in melanic scales, and a possible but poorly understood interaction with the ommochrome pathway in red scales. Scale bars : D, H = 100 µm.

**Figure S9.**
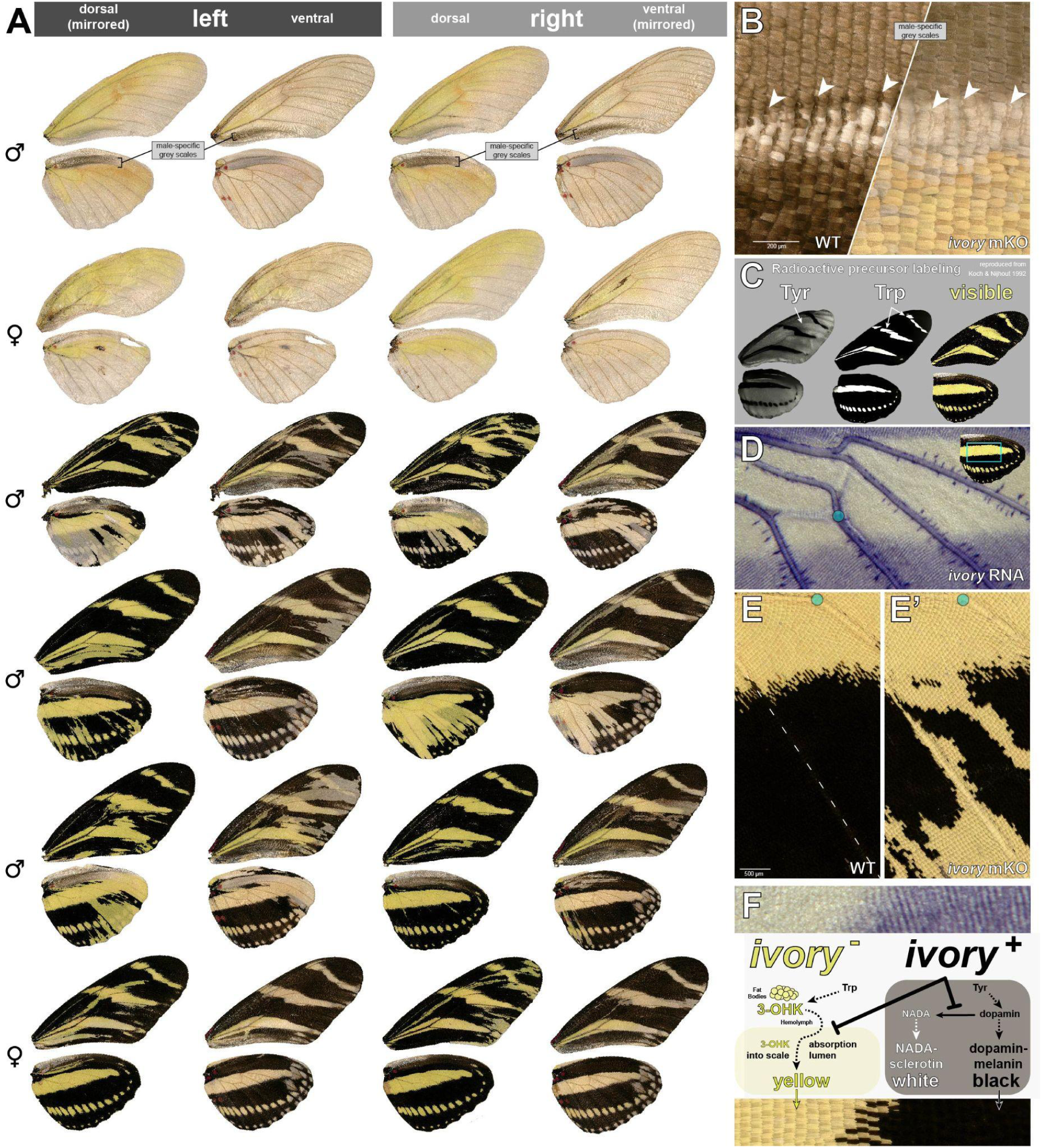
Mosaic knock-outs of *ivory* suggest a dual role in *H. charithonia*. **(A)** Representative *ivory* crispant phenotypes in *H. charithonia*. Mutant clones show a yellow colouration, most pronounced on dorsal sides. Male crispants show a residual patch of grey scales in the wing-apposition region (anal region of ventral forewings, anterior region of dorsal hindwings), that is not observed in females. **(B)** Magnified view of the WT *vs. ivory* mKO in the wing apposition, here centred on the Rs vein of dorsal hindwings (horizontal line of white scales), which bears the male androconial scales (arrowheads). Androconial scales are depigmented in *ivory* crispants. **(C)** Reproduction of wing autoradiographs (56) marking the incorporation of isotope-labeled Tyrosine (marking presumptive melanins), and Tryptophan (marking ommochrome-pigment regions). Autoradiograph images from the original publication were re-processed with Denoising tools in Adobe Photoshop, and are meant to convey the original results schematically here. 3-OHK is known to be the yellow ommochrome pigment in this species (57). Melanins and 3-OHK are spatially exclusive in WT butterflies, implying that melanins do not mask a broadly distributed 3-OHK. **(D)** *In situ* hybridization of *ivory* around 35% of pupal development in the dorsal hindwing bar region. Cyan : M_3_-crossvein intersection. **(E-E’)** Example of ectopic yellow colouration in ivory mutant clones, here in the region distal to the yellow hindwing bar. **(F)** Working model for explaining the yellow colouration of *ivory*-deficient clones. In addition to fostering dopamine-melanin scale fate (black scale), *ivory* necessarily prevents the incorporation of 3-OHK in these scales (panel C).

**Figure S10.**
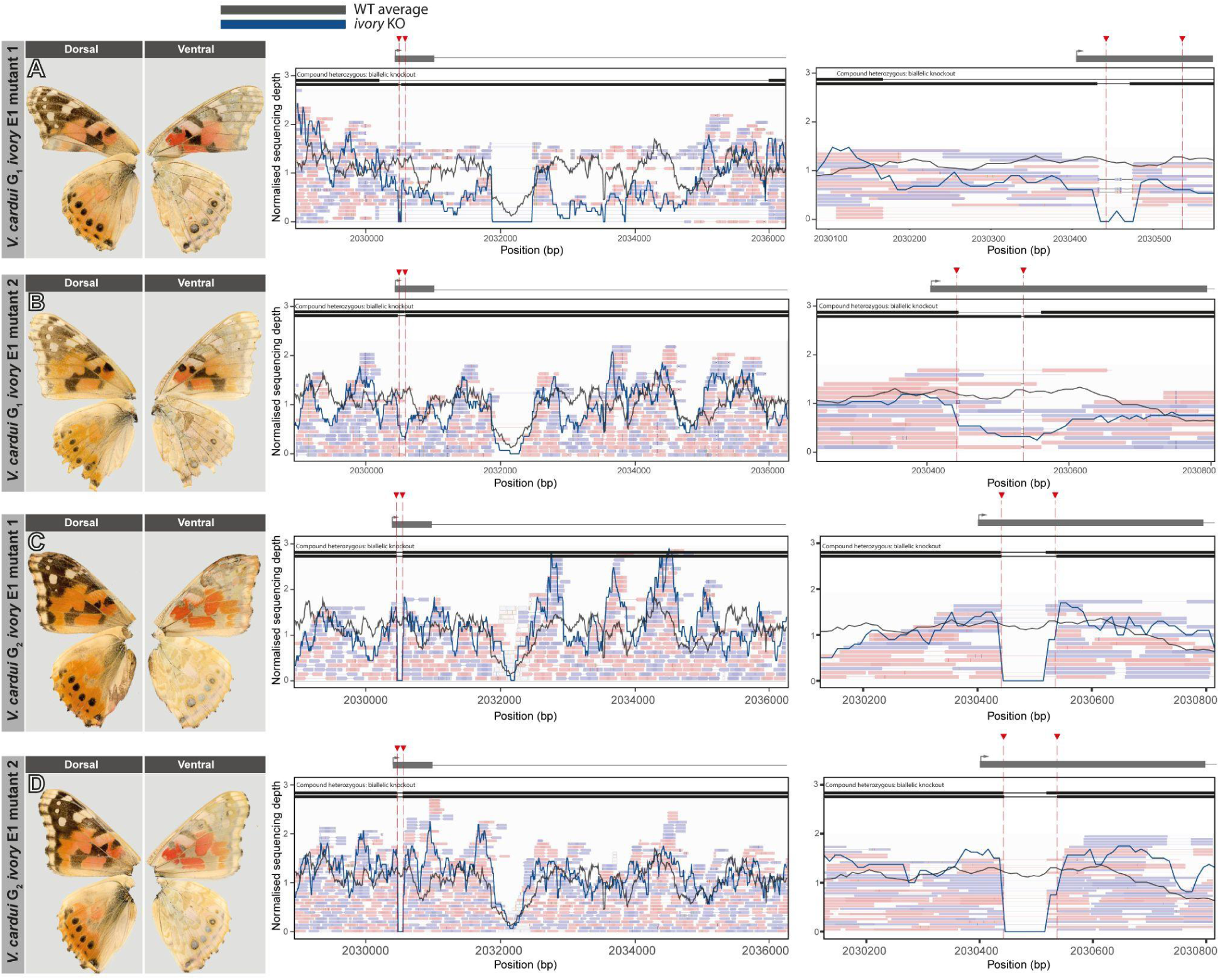
Genotyping of G_1_ and G_2_ *ivory exon 1* mutants. **(A)** Partly depigmented G_1_ *ivory* mutant displaying a compound heterozygous mutation. One allele carries a 32 bp mutation centred around the left cut-site. A second allele, partially overlapping with the first, carries a 6.14 kb deletion encompassing all of exon 1 and part of the first intron of *ivory*. **(B)** Partly depigmented G_1_ *ivory* mutant displaying a smaller compound heterozygous mutation, with one small deletion allele of 4 bp and a larger partially overlapping allele with a 121 bp deletion. **(C)** Strongly depigmented G_2_ *ivory* mutant, displaying a compound heterozygous KO. Two deletion alleles of similar size (82 bp and 97 bp) and overlapping over 78 bp are shown. **(D)** Replicate sample from the same rearing batch, genotyped for the same deletion alleles as in panel C.

**Figure S11.**
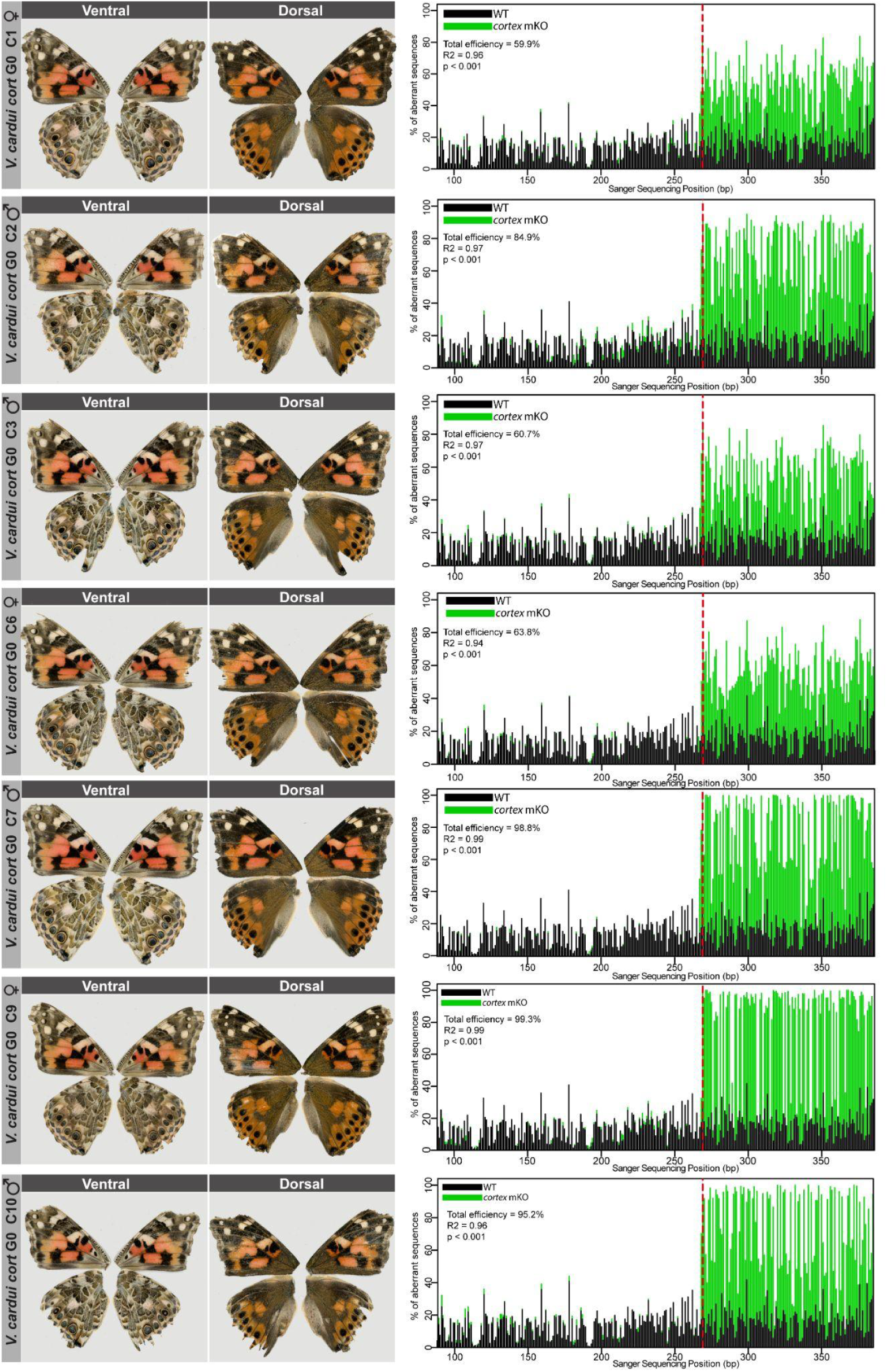
Genotyping of *cortex exon 2* G_0_ mutants. Individuals displaying *cortex* mutations are shown on the left, both dorsal and ventral views are displayed. TIDE analysis (80) next to each individual is shown on the right, indicating evidence of mutations close to the expected sgRNA cure site (dashed red line). Calculated total efficiency is based on % mutant signal from chromatogram decomposition and associated correlation and significance are shown on each plot.

**Figure S12.**
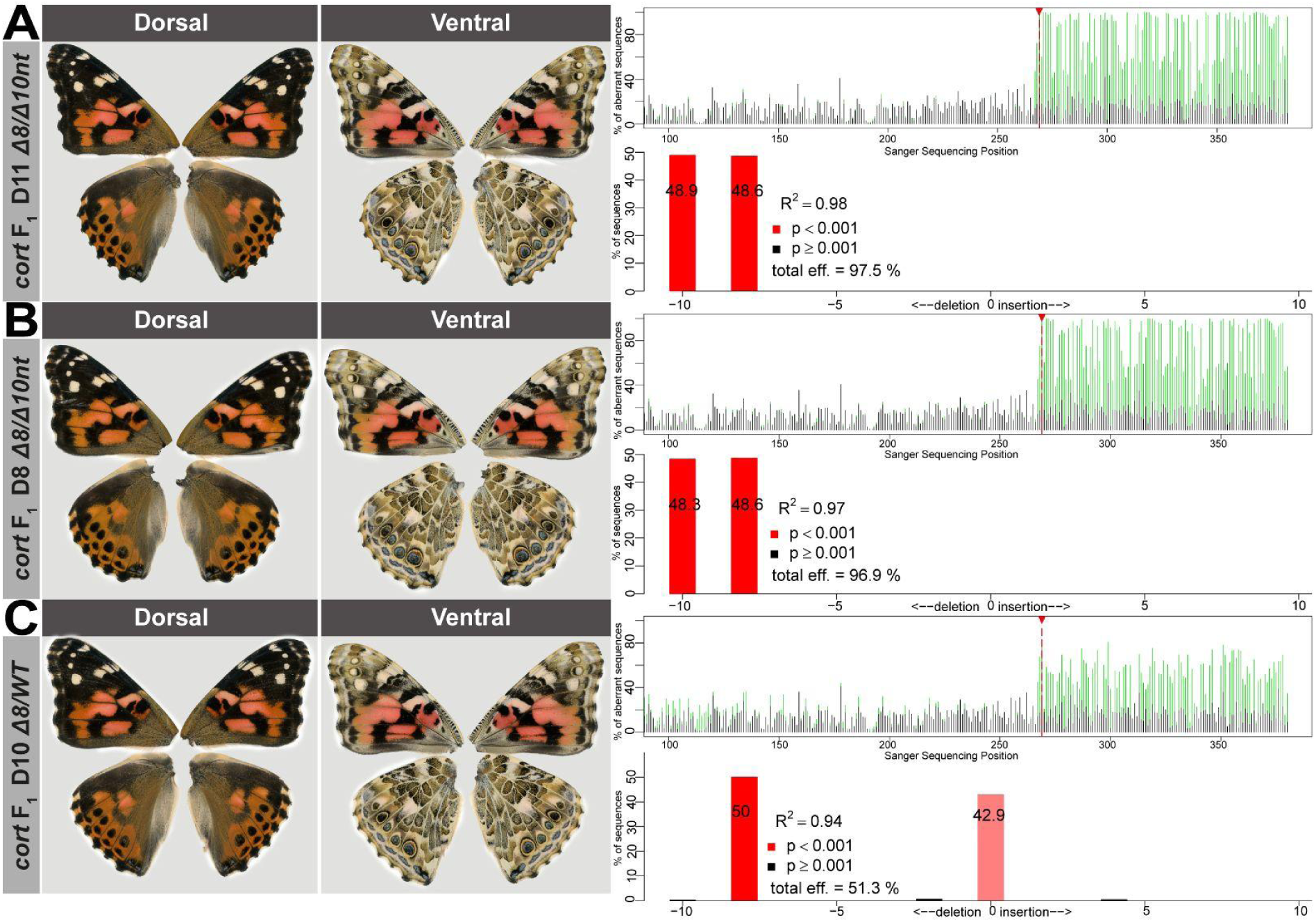
Genotyping of *cortex exon 2* F_1_ mutants. **(A-B)** Examples of compound heterozygous mutants displaying a 8 nt and 10 nt deletion, as shown by TIDE analysis of Sanger sequence chromatograms at the expected CRISPR cuting site (red dotted line). **(C)** A true heterozygous mutant carrying an 8 nt deletion and a WT allele. No wing phenotypes are detectable in these mutants.

**Figure S13.**
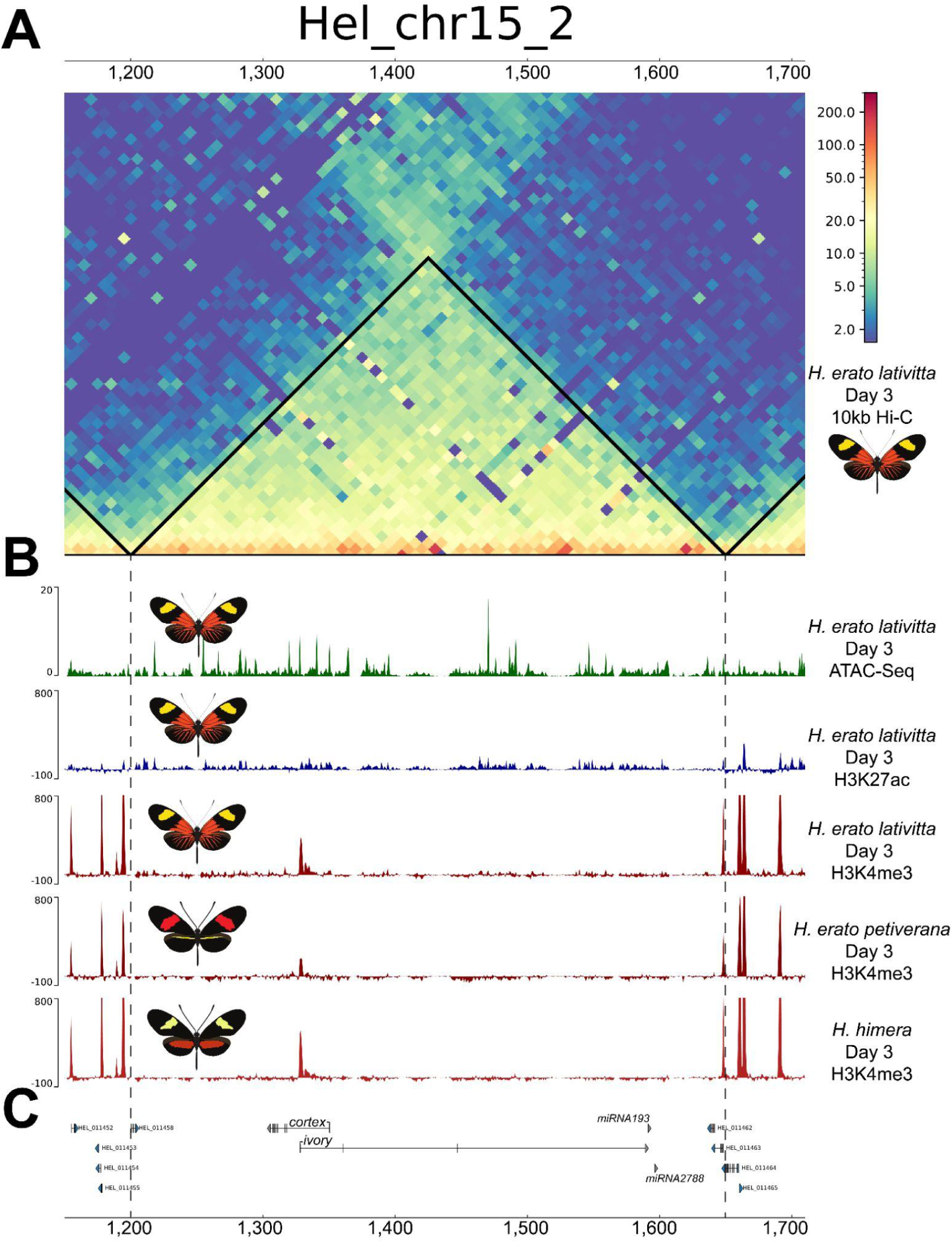
The *ivory* promoter occurs within a TAD with no other sites of transcriptional poising. **(A)** Hi-C contact heatmap in Day 3 pupal forewings of *H. erato lativitta* at 10-kb binning. A TAD encompasses both *ivory* and *cortex* transcripts (boundaries indicated by vertical dotted lines). **(B)** Chromatin accessibility, H3K27ac and H3K4me3 ChIPseq enrichment profiles are shown for *H. erato lativitta*. The unique H3K4me3 peak within the TAD indicates transcriptional poising and activity of the *ivory* promoter. This H3K4me3 mark is also present in *H. erato petiverana* and *H. himera*. **(C)** Annotation of *ivory* and *cortex* in *H. erato petiverana*; *miR193* and *miR2788* are shown at the 3’ end of *ivory*.

**Figure S14.**
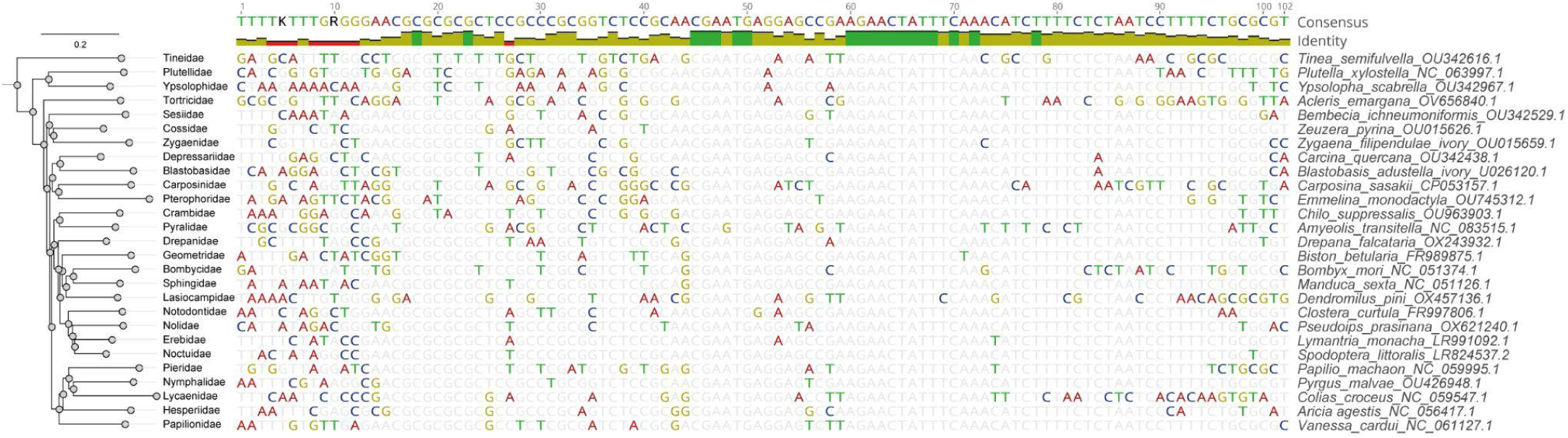
Multiple sequence alignment showing the putative *ivory* promoter across the Dytrisia lineage. Recovered BLAST hits from representative species from each lepidopteran family are shown as a multiple alignment spanning a ∼100 bp region of high conservation. GenBank accession numbers of the extracted scaffold containing the *ivory* locus are indicated for each species. Phylogeny is based on ((89)).

**Figure S15.**
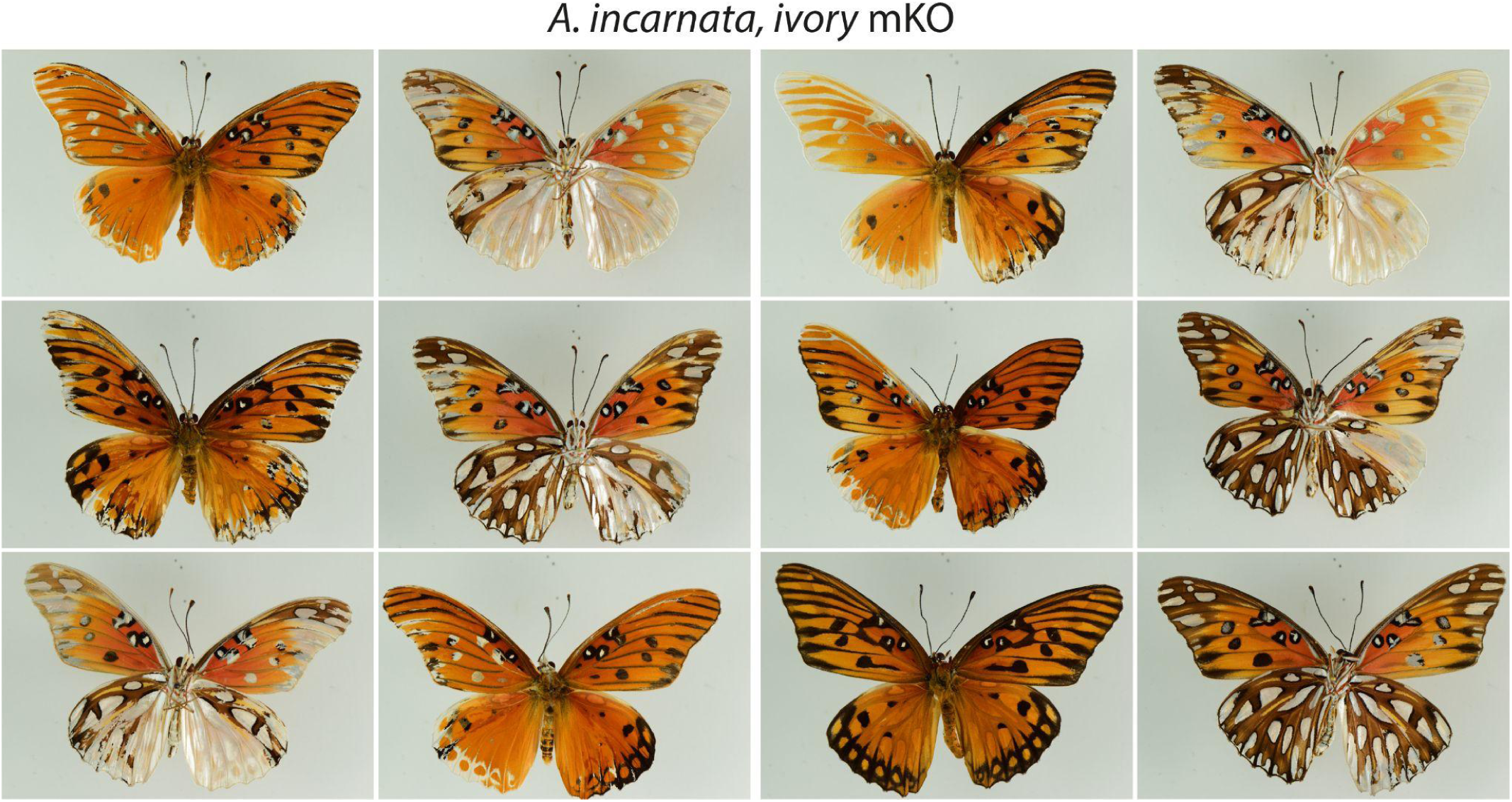
Representative *A. incarnata ivory* crispant phenotypes. Ventral (left image) juxtaposed to dorsal (right image) sides for each individual.

**Figure S16.**
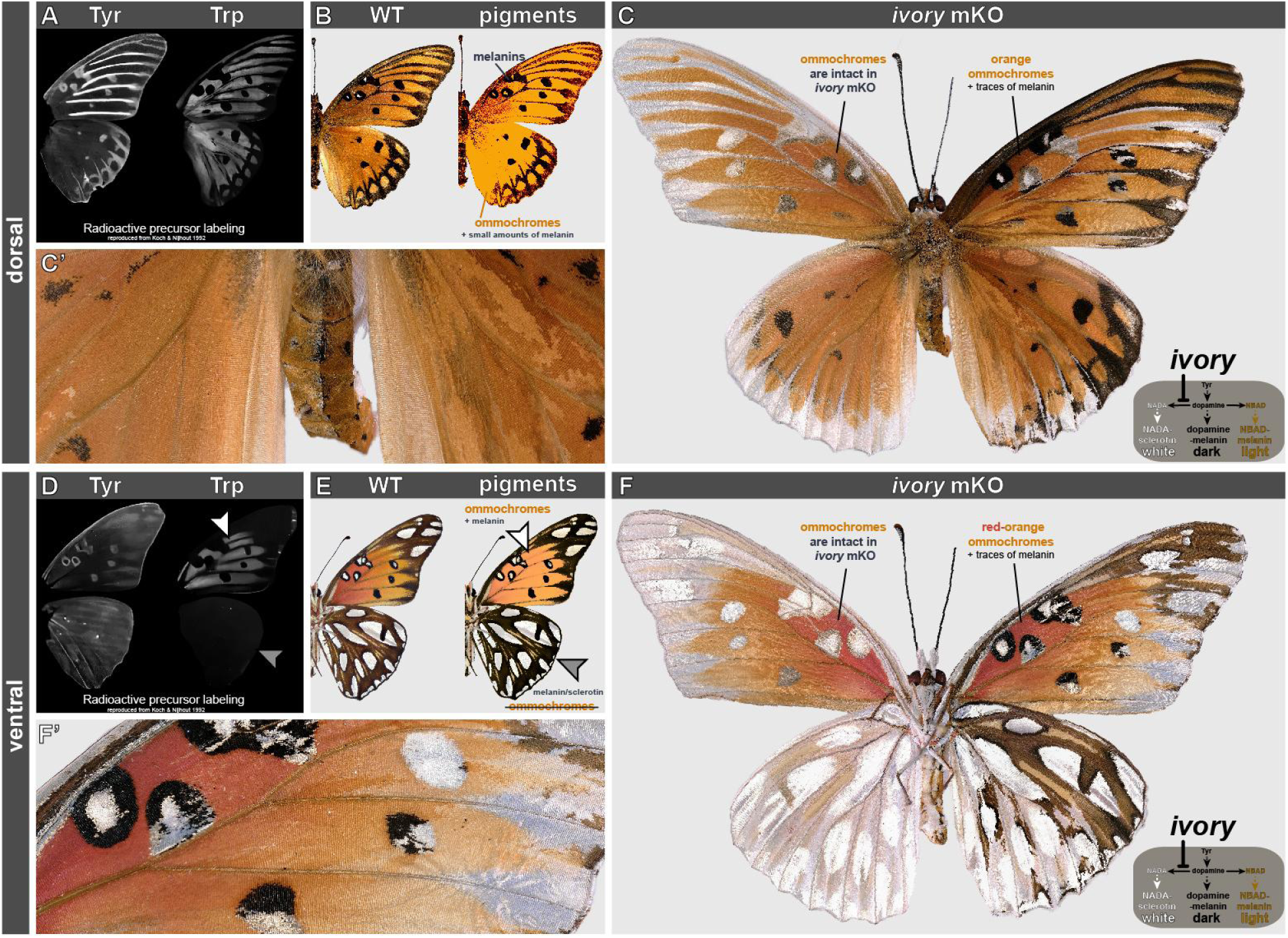
Multiple effects of *ivory* mKO on *A. incarnata* colour patterns. **(A)** Autoradiographs for tyrosine and tryptophan incorporation in dorsal wing surfaces, reproduced from a previous publication (56). Tyrosine is ubiquitous, but strongly marking presumptive black patterns. Tryptophan is limited to presumptive orange patterns. **(B)** Resulting model for melanin (Tyr-metabolism) vs ommochrome (Trp-metabolism) on *A. incarnata* dorsal wings. **(C-C’)** Example of *A. incarnata* G_0_ female crispant for *ivory*. The left side of this butterfly appears to be mosaic-mutant (see midline split over the abdomen, magnified in C’), while the right side is mosaic. Black-to-silver transformations are consistent with a dopamine-melanin (dark melanin) to NADA-sclerotin (colourless/white) conversion. Lighter orange clones are consistent with the presence of melanin species, lost or converted to clear in the mKO. (**D-F’**) Same as (A-D’) but on ventral surfaces. Tryptophan incorporation is specific to the ventral forewing and absent from the ventral hindwing (E, arrowheads). Accordingly, red-orange ventral forewing pigments are ommochromes, and persist in the *ivory* mKO, albeit a lighter appearance (magnified in D’), due to the likely admixture of melanins (faint Tyr signal in D). C and F show the same crispant specimen, with F mirror-reversed to facilitate dorso-ventral comparisons.

**Figure S17.**
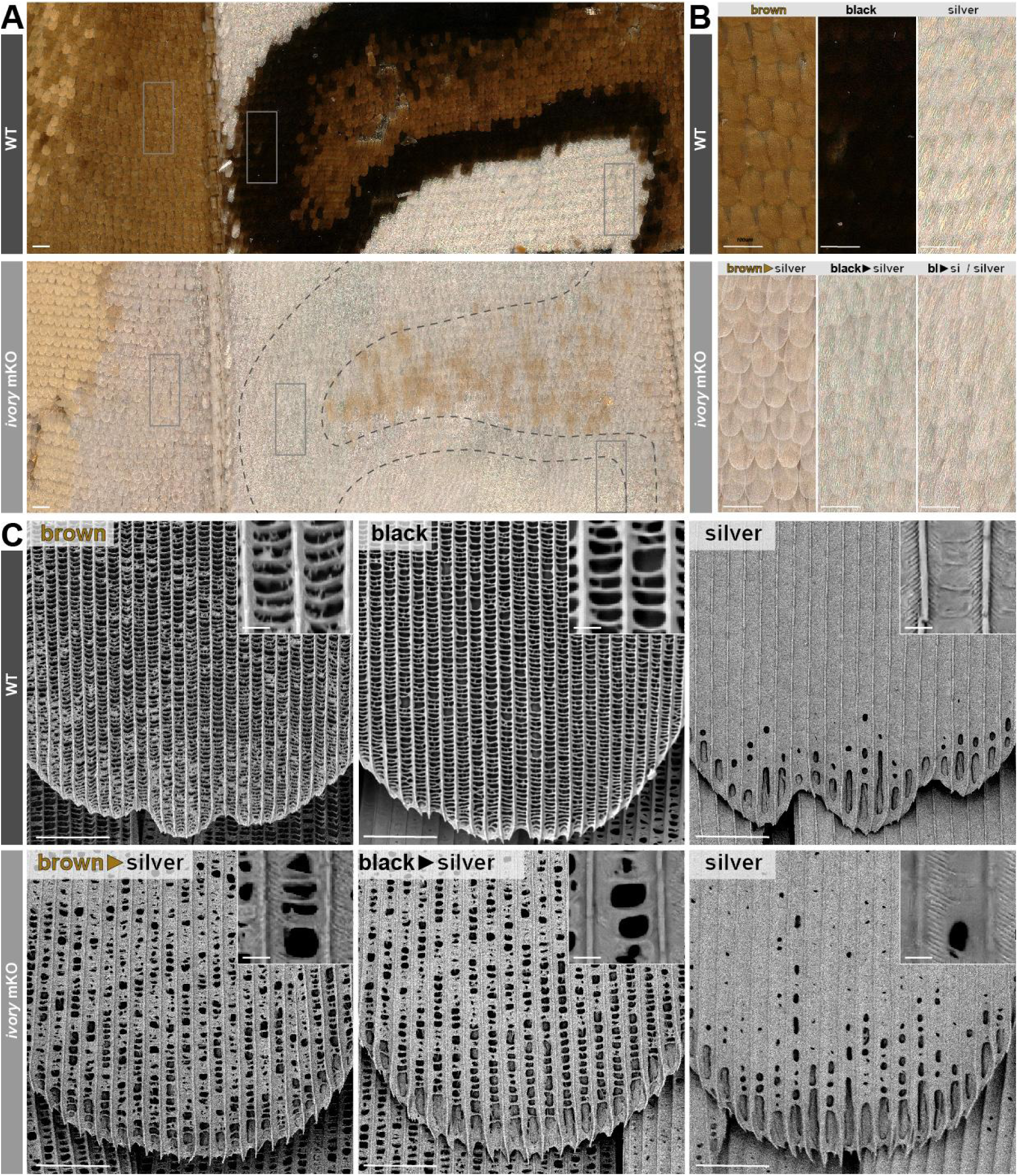
Ultrastructural effects of *ivory* mKO in *A. incarnata* transformed scales. **(A-B).** Magnified views of the ventral anterior hindwings of *A. incarnata*, comparing WT and *ivory* mKO conditions. Insets (rectangles) highlight colour conversions from WT to crispant states in presumptive brown (leftmost), black (center, within dashed lines), and silver (rightmost) regions. Presumptive brown scales show variable conversion phenotypes, ranging from silver, to white (less specular, shown in B), to light brown. **(C)** Scanning electron micrographs of representative scales in the two conditions. Transformed melanic scales show ectopic lamination partially covering the upper scale surface, resembling the WT condition. Scale bars : A-B = 100 µm; C = 10 µm; C, insets = 1 µm.

**Figure S18.**
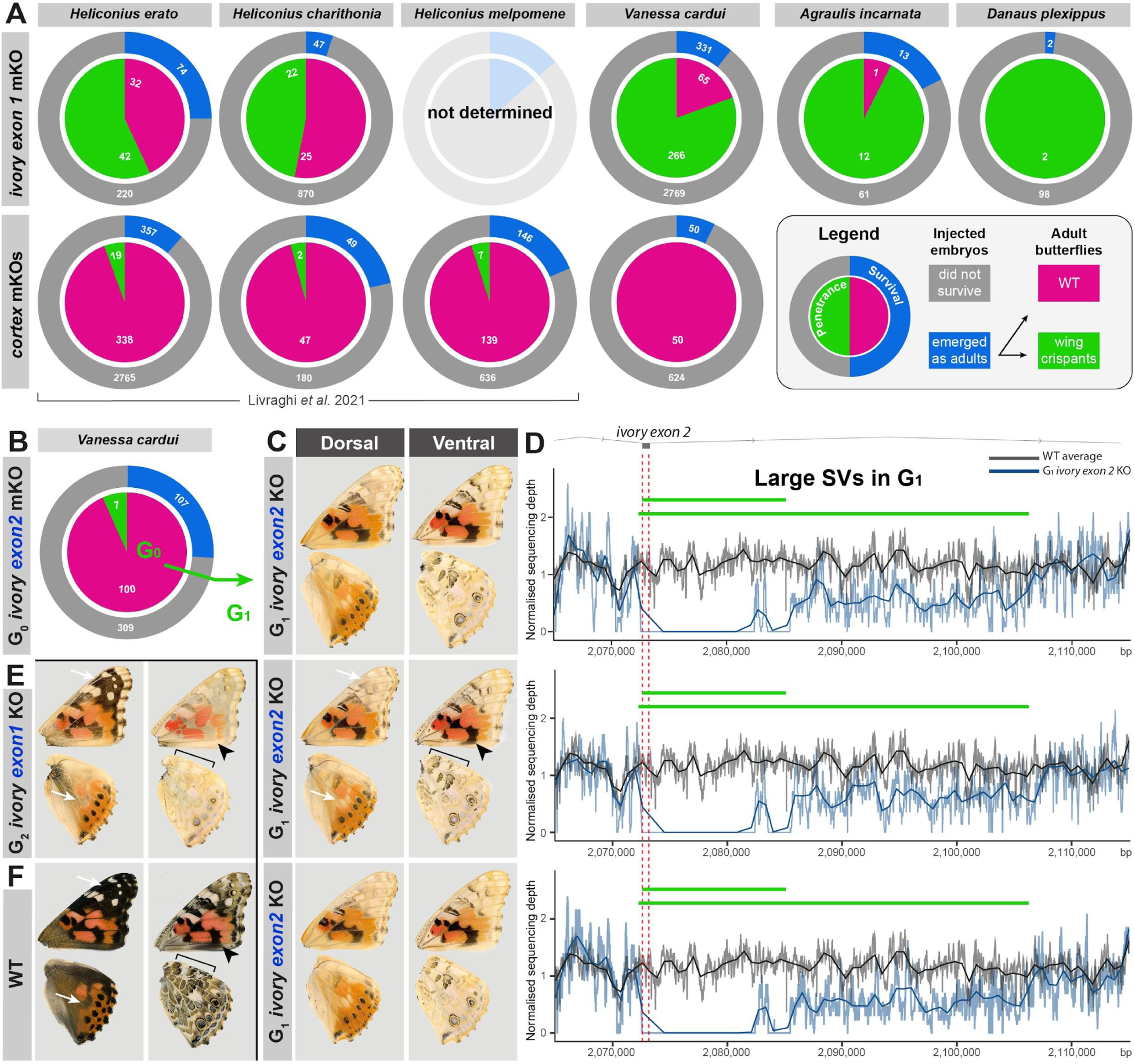
**(A)** Multiple species comparison of CRISPR G_0_ colour phenotypes, based on Table S1 and a previous report for *cortex* mKO experiments (Livraghi et al. 2021). Outer donut charts show the proportion of injected embryos resulting in surviving adults (blue). Inner pie charts show the proportion of emerging adults carrying any wing phenotype (green). **(B)** Low penetrance of *V. cardui* G_0_ colour phenotypes when targeting *ivory exon 2*. **(C-D)** Partial colour phenotype and genotyping-by-sequencing of *V. cardui* G_1_ mutant siblings, following the targeting of *ivory exon 2*. Coverage analysis shows the three individuals are compound heterozygotes for two unexpectedly large deletions (green bars), extending from *exon 2* (red lines = sgRNA targets) into the second intron of *ivory*. Wing phenotypes are nearly identical among the three siblings. **(E-F)** Wing phenotypes from a confirmed G_2_ *ivory exon 1* compound heterozygote (**Fig. S11D**) and *V. cardui* WT individual. Compared to *exon1* mutants, the large SV compound heterozygotes (C-D) show hypomorphic effects, as seen on central forewing black patterns (black arrowhead) and on the ventral hindwing Central Symmetry System (bracket), but also a strong depigmentation phenotype on dorsal distal forewings and the median dorsal hindwing (white arrows).

### Supporting Text

#### CRISPR mutagenesis can induce fortuitous large SVs that associate with low-penetrance G_0_ crispant phenotypes

CRISPR targeting of the *ivory promoter/exon 1* provided adult G_0_ phenotypes with a high penetrance range of 45-100% across 5 species in this study (**Fig. S18A**). To probe the functionality of downstream elements in this gene, we mutagenized the *exon 2* of *ivory* in *V. cardui,* an element isolated from 3’RACE in this species but that is not shared or alignable with *ivory* sequences in *Heliconius spp*. These mosaic knock-outs resulted in partial pigmentation phenotypes, with a low penetrance of 7%, instead of 80% when targeting *exon 1* in this species (**Fig. S18B**). Crossing two of the G_0_ crispants propagated pigmentation defects into a G_1_ generation, enabling proper genotyping (**Fig. S18C,D**). To our surprise, aligning short-read sequencing data from a total of three *ivory exon 2* G_1_ mutant siblings revealed compound heterozygosity for two large structural variants (SVs). Specifically, coverage drop-out near the CRISPR targeted region indicates a > 30-kb long deletion extending into the second intron of *ivory,* as well as a second deletion of > 12 kb. These two deletions overlap over a 12.9 kb interval with no read-mapping, with the exception of a site with inferred transposable elements. Thus, while low-penetrance in this experiment suggests the non-conserved *exon 2* of the *V. cardui ivory* lncRNA is facultative or non-functional, we nonetheless obtained G_0_ phenotypes by sporadically generating large SVs at this locus. In summary, we have circumstantial evidence that low-penetrance phenotypes can result from large SVs induced by CRISPR targeted mutagenesis.

We formally showed in this study that *cortex* has no colour patterning function, at least in *V. cardui*. This somewhat contradicts previous *cortex* CRISPR experiments in *Heliconius spp.,* which obtained G_0_ phenotypes (6), albeit with small clones and an adult penetrance of only 4-5% (*ivory* mKOs showed a penetrance of 45-56% in these species). To explain these reported phenotypes and their low penetrance altogether, we propose that *cortex* mKO effects were the result of unintendedly large SVs interfering with *ivory* expression. Proving this directly in *Heliconius* would be challenging, as it would imply re-generating and propagating sporadic mutations into a live G_1_ or G_2_ generation before genome resequencing. Nonetheless, *ivory* targeting experiments provided a ten-fold increase in two *Heliconius* species and produced with large crispant clones, in line with the expectations of a high-efficient CRISPR target at a non-lethal locus. In summary, comparison of all CRISPR experiments strongly suggests that *cortex* targeting produced fortuitous ivory-deficient alleles, a phenomenon that generally urges caution in the interpretation of low-penetrance crispant data in the future.

#### Full length ivory transcripts recovered from *Heliconius* Trinity assemblies

##### Heliconius erato

>Ivory_Iso1_TRINITY_DN20323_c0_g1_i1 [organism=Heliconius erato] ivory isoform assembled from wing transcriptome

CTCATTCGTCGCGTAGATCGCGAGCTGCGCGCGCGCGTACCGCACACATTTAGTCACGCTTGATCGCCAGGGCAACGACCTCGCGGCCATTTTTTTCCAACGAACGTCCTTGCCAAACATGGCGAACAGCTGATGCACGATCCAATCTGTGTAATTTTTAAGTGGCTCTATCACTTGTAACTATTTAAATAAAAATTAACATTTTTTCTTGTCAGTACTGGCCAGTATATATATATTTTTTATTTATTAAAATTAAAATAGTGACAAATTAGTGTGGTGTGGTGCCTTATATTGTAATTTGATTGAATGTGATGTGTTTGTGCCTGTGATGATAAGATTATTTATGTTACAGGCAAAGAAGCGTCGCTGTGGCATCGACAGTGCGTCAATATAGTTTGGTCCCCGGCACGGCATCGGTATCAAATTATAAGCAAGGTGTTATTAGAGTTCAATTATTTTTGCAAGATAGGAGGTAGACGTGTAAATCGAAGATTCGAACTGGAAAGAAGTATTAAGAAGTTGAGGACAGTGGACGAGACGAACGACTTAAGCAGAGNCCGTAGCGTTTGGCTCTCCTTTCTCTCTGGCCACTCCGCTAGTGACAATGCGTTAAGATATCACGAGAACTAAATGCTGGATGACCTATAGCTGGGTAACATTTTAAAAAGAAAGTTTATAAAATATTATATTTACTTAGTAAAATAGAAATAAATTAATGTTATCAGTTAACGTGGAATAGTGAACTTTGCTAGAGTGCACGCTATAACGGACGTTCCGCAAAATCGTCATGGCAAATTTAAAGGCATAATAAAGAATTATTTTTGTTAAATATATTTTTTTCTATTTGATTTTTTGTATTTAAGCCCTGTGCGACTACGCCCG

>Ivory_Iso2_TRINITY_DN20323_c0_g1_i5 [organism=Heliconius erato] ivory isoform assembled from wing transcriptome

CTCATTCGTCGCGTAGATCGCGAGCTGCGCGCGCGCGTACCGCACACATTTAGTCACGCTTGATCGCCAGGGCAACGACCTCGCGGCCATTTTTTTCCAACGAACGTCCTTGCCAAACATGGCGAACAGCTGATGCACGATCCAATCTGTGTAATTTTTAAGTGGCTCTATCACTTGTAACTATTTAAATAAAAATTAACATTTTTTCTTGTCAGTACTGGCCAGTATATATATATTTTTTATTTATTAAAATTAAAATAGTGACAAATTAGTGTGGTGTGGTGCCTTATATTGTAATTTGATTGAATGTGATGTGTTTGTGCCTGTGATGATAAGATTATTTATGTTACAGGCAAAGAAGCGTCGCTGTGGCATCGACAGTGCGTCAATATAGTTTGGTCCCCGGCACGGCATCGAAAACTTGGTGGCCCTACTAAGTTGGCAAATATCCATCACGCCGCACGATACGCTAGAAATATGTATCAAATTATAAGCAAGGTGTTATTAGAGTTCAATTATTTTTGCAAGATAGGAGGTAGACGTGTAAATCGAAGATTCGAACTGGAAAGAAGTATTAAGAAGTTGAGGACAGTGGACGAGACGAACGACTTAAGCAGAGNCCGTAGCGTTTGGCTCTCCTTTCTCTCTGGCCACTCCGCTAGTGACAATGCGTTAAGATATCACGAGAACTAAATGCTGGATGACCTATAGCTGGGTAACATTTTAAAAAGAAAGTTTATAAAATATTATATTTACTTAGTAAAATAGAAATAAATTAATGTTATCAGTTAACGTGGAATAGTGAACTTTGCTAGAGTGCACGCTATAACGGACGTTCCGCAAAATCGTCATGGCAAATTTAAAGGCATAATAAAGAATTATTTTTGTTAAATATATTTTTTTCTATTTGATTTTTTGTATTTAAGCCCTGTGCGACTACGCCCG

>Ivory_Iso3_TRINITY_DN20323_c0_g1_i6 [organism=Heliconius erato] ivory isoform assembled from wing transcriptome

CTCATTCGTCGCGTAGATCGCGAGCTGCGCGCGCGCGTACCGCACACATTTAGTCACGCTTGATCGCCAGGGCAACGACCTCGCGGCCATTTTTTTCCAACGAACGTCCTTGCCAAACATGGCGAACAGCTGATGCACGATCCAATCTGTGTAATTTTTAAGTGGCTCTATCACTTGTAACTATTTAAATAAAAATTAACATTTTTTCTTGTCAGTACTGGCCAGTATATATATATTTTTTATTTATTAAAATTAAAATAGTGACAAATTAGTGTGGTGTGGTGCCTTATATTGTAATTTGATTGAATGTGATGTGTTTGTGCCTGTGATGATAAGATTATTTATGTTACAGGCAAAGAAGCGTCGCTGTGGCATCGACAGTGCGTCAATATAGTTTGGTATCAAATTATAAGCAAGGTGTTATTAGAGTTCAATTATTTTTGCAAGATAGGAGGTAGACGTGTAAATCGAAGATTCGAACTGGAAAGAAGTATTAAGAAGTTGAGGACAGTGGACGAGACGAACGACTTAAGCAGAGNCCGTAGCGTTTGGCTCTCCTTTCTCTCTGGCCACTCCGCTAGTGACAATGCGTTAAGATATCACGAGAACTAAATGCTGGATGACCTATAGCTGGGTAACATTTTAAAAAGAAAGTTTATAAAATATTATATTTACTTAGTAAAATAGAAATAAATTAATGTTATCAGTTAACGTGGAATAGTGAACTTTGCTAGAGTGCACGCTATAACGGACGTTCCGCAAAATCGTCATGGCAAATTTAAAGGCATAATAAAGAATTATTTTTGTTAAATATATTTTTTTCTATTTGATTTTTTGTATTTAAGCCCTGTGCGACTACGCCCG

>Ivory_Iso4_TRINITY_DN20323_c0_g1_i8 [organism=Heliconius erato] ivory isoform assembled from wing transcriptome

CTCATTCGTCGCGTAGATCGCGAGCTGCGCGCGCGCGTACCGCACACATTTAGTCACGCTTGATCGCCAGGGCAACGACCTCGCGGCCATTTTTTTCCAACGAACGTCCTTGCCAAACATGGCGAACAGCTGATGCACGATCCAATCTGTGTAATTTTTAAGTGGCTCTATCACTTGTAACTATTTAAATAAAAATTAACATTTTTTCTTGTCAGTACTGGCCAGTATATATATATTTTTTATTTATTAAAATTAAAATAGTGACAAATTAGTGTGGTGTGGTGCCTTATATTGTAATTTGATTGAATGTGATGTGTTTGTGCCTGTGATGATAAGATTATTTATGTTACAGGCAAAGAAGCGTCGCTGTGGCATCGACAGTGCGTCAATATAGTTTGAAAACTTGGTGGCCCTACTAAGTTGGCAAATATCCATCACGCCGCACGATACGCTAGAAATATGTATCAAATTATAAGCAAGGTGTTATTAGAGTTCAATTATTTTTGCAAGATAGGAGGTAGACGTGTAAATCGAAGATTCGAACTGGAAAGAAGTATTAAGAAGTTGAGGACAGTGGACGAGACGAACGACTTAAGCAGAGNCCGTAGCGTTTGGCTCTCCTTTCTCTCTGGCCACTCCGCTAGTGACAATGCGTTAAGATATCACGAGAACTAAATGCTGGATGACCTATAGCTGGGTAACATTTTAAAAAGAAAGTTTATAAAATATTATATTTACTTAGTAAAATAGAAATAAATTAATGTTATCAGTTAACGTGGAATAGTGAACTTTGCTAGAGTGCACGCTATAACGGACGTTCCGCAAAATCGTCATGGCAAATTTAAAGGCATAATAAAGAATTATTTTTGTTAAATATATTTTTTTCTATTTGATTTTTTGTATTTAAGCCCTGTGCGACTACGCCCG

##### Heliconius melpomene

>Ivory_Iso1_TRINITY_DN1775_c7_g1_i6 [organism=Heliconius melpomene] ivory isoform assembled from wing transcriptome

AAACAATTTTTATTGTCAGTACTGGCCAGTATATATTTTTATTTACTTTTTTTATTTATTAAAATTAAAACAGTGACAAATTAGTGTGGTGTGGTGCCTTATATTATTGATTAAATGTGATGTGTTTGTGCCTGTGATGATAACATTATTTATGTTATAGCGAAGAACGCGTCGCTGTGACACTGACAGTGCGTCAATATAGTTTGAAAACTTGGTGGCCCTACTAACTTGGCAAATATCCATCACGCCGCACGATACGCAACAAATATGTATCAAATTCTGAGCTACAGTTCAATTATATCACGAGAACTAAACGCTGGATGACCTATAGCTGGGTAACATTTTAAAAAGAAAGTCCATAAAATATTATATTTACTGTGGAATTCAATTAATGCTATCAGATAACGTGGAAAAATGAACTTTGCTAGAGTGCACGCTATAACAGTTGTTCCACAAAATGAATGGCAAATTTTAAGGCGTAATAAAGAATTTTAGTTTTTTCTAAAGTATTTTTTACATATTTGAATTTTTGAATTTTAAGTTTAATCTTTTGTGATTTATAAATCACAAAAAAATATTGTACTTTCGGTATAATTTAAATATTTGTTTTTCTTAACTTCTTTTATTTATAGAATTATCTTTTATAAAAAGGATATTTTATTACGTAGCCCGTAGCTTTTACATATATACATTACATTATATATCTATATTAATACTTGAAGCAAAAATTTGGTACCTCGATAAAATTGAGGGTGGGGAACGTAGTGTCTTAACACTCGGTTCCCTCTGCCTACCCTGCTGGGTGCGGGATACAGCGTGAAGCTGATTTATTTATTTGATTTATTTGATAAAATTGGGCACAGAGAAGTAGAAA

>Ivory_Iso3_TRINITY_DN1319_c0_g1_i28 [organism=Heliconius melpomene] ivory isoform assembled from wing transcriptome

TTTTATTTACTTTTTTTATTTATTAAAATTAAAACAGTGACAAATTAGTGTGGTGTGGTGCCTTATATTATTGATTAAATGTGATGTGTTTGTGCCTGTGATGATAACATCATTTATGTTATAGCAAAGAACGCGTCGCTGTGACACTGACAGTGCGTCAATATAGTTTGGTCCACGGCACGGCATCGAAAACTTGGTGGCCCTACTAACTTGGCAAATATCCATCACGCCGCACGATACGCAACAAATATGTATCAAATTCTGAGCTACAGTTCAATTATATCACGAGAACTAAACGCTGGATGACCTATAGCTGGGTAACATTTTAAAAAGAAAGTCCATAAAATATTATATTTACTGTGGAATTCAATTAATGCTATCAGATAACGTGGAAAAATGAACTTTGCTAGAGTGCACGCTATAACAGTTGTTCCACAAAATGAATGGCAAATTTTAAGGCGTAATAAAGAATTTTAGTTTTTTCTAAAGTATTTTTTAAATATTTGAAGTTTTGTATTTTAAACTTAAAATATCTGTGATT

>Ivory_Iso4_TRINITY_DN1319_c0_g1_i27 [organism=Heliconius melpomene] ivory isoform assembled from wing transcriptome

TTTTATTTACTTTTTTTATTTATTAAAATTAAAACAGTGACAAATTAGTGTGGTGTGGTGCCTTATATTATTGATTAAATGTGATGTGTTTGTGCCTGTGATGATAACATCATTTATGTTATAGCAAAGAACGCGTCGCTGTGACACTGACAGTGCGTCAATATAGTTTGAAAACTTGGTGGCCCTACTAACTTGGCAAATATCCATCACGCCGCACGATACGCAACAAATATGTATCAAATTCTGAGCTACAGTTCAATTATATCACGAGAACTAAACGCTGGATGACCTATAGCTGGGTAACATTTTAAAAAGAAAGTCCATAAAATATTATATTTACTGTGGAATTCAATTAATGCTATCAGATAACGTGGAAAAATGAACTTTGCTAGAGTGCACGCTATAACAGTTGTTCCACAAAATGAATGGCAAATTTTAAGGCGTAATAAAGAATTTTAGTTTTTTCTAAAGTATTTTTTAAATATTTGAAGTTTTGTATTTTAAACTTAAAATATCTGTGATT

>Ivory_Iso5_TRINITY_DN1319_c0_g1_i11 [organism=Heliconius melpomene] ivory isoform assembled from wing transcriptome

TTTTATTTACTTTTTTTATTTATTAAAATTAAAACAGTGACAAATTAGTGTGGTGTGGTGCCTTATATTATTGATTAAATGTGATGTGTTTGTGCCTGTGATGATAACATCATTTATGTTATAGAAAACTTGGTGGCCCTACTAACTTGGCAAATATCCATCACGCCGCACGATACGCAACAAATATGTATCAAATTCTGAGTACAGTTCAATTATATCACGAGAACTAAACGCTGGATGACCTATAGCTGGGTAACATTTTAAAAAGAAAGTCCATAAAATATTATATTTACTGTGGAATTCAATTAATGCTATCAGATAACGTGGAAAAATGAACTTTGCTAGAGTGCACGCTATAACAGTTGTTCCACAAAATGAATGGCAAATTTTAAGGCGTAATAAAGAATTTTAGTTTTTTCTAAAGTATTTTTTAAATATTTGAAGTTTTGTATTTTAAACTTAAAATATCTGTGATT

### Supplementary Tables

**Table S1.** Summary of CRISPR injection experiments.

| Species | sgRNA(s) | Inj. Embryos Ninj | L1 larvae Nhat | Pupae | Adults Nadu | Crispants Nmut | Inj. time h AEL | Cas9:sgRNA ng/μL | Hatching Rate Nhat/Ninj | Penetrance Nmut/Ninj |
| --- | --- | --- | --- | --- | --- | --- | --- | --- | --- | --- |
| *H. erato* | *He_ivory12* | 64 | 51 | 24 | 18 | 12 | 1-4 | 500:125:125 | 79.7% | 66.7% |
|  |  | 230 | 172 | 80 | 56 | 30 | 1-2 | 500:125:125 | 74.8% | 53.6% |
| *H. charithonia* | *He_ivory1+2* | 917 | 441 | 73 | 47 | 22 | 0-1 | 500:250:250 | 48.1% | 46.8% |
| *A. incarnata* | *Ai_ivory1+2* | 74 | 36 | 16 | 13 | 12 | 0.5-2 | 500:250 | 48.6% | 92.3% |
| *D. plexippus* | *Dp_ivory1+2+3* | 100 | 24 | 7 | 2 | 2 | 0-1 | 500:166(each) | 24.0% | 100.0% |
| *V. cardui* | *Vc_ivory1+2* | 1113 | 149 | 220 | 49 | 41 | 0.75-1.5 | 500:125:125 | 13.4% | 83.7% |
| *V. cardui* | *Vc_ivory1+2* | 1250 | 174 | 382 | 65 | 56 | 2-3 | 500:125:125 | 13.9% | 86.2% |
| *V. cardui* | *Vc_ivory1+2* | 737 | n.d. | n.d. | 217 | 169 | 0.75-2 | 500:125:125 | n.d. | 77.9% |
| *V. cardui* | *Vc_ivoryE2_1+2* | 416 | 232 | 116 | 107 | 7 | 2.5-3.5 | 500:250 | 55.8% | 6.5% |
| *V. cardui* | *Vc_cortex* | 674 | 370 | 53 | 50 | 0 | 1.5-2 | 500:250 | 54.9% | 0.0% |

**Table S2.** Comparison of survival and penetrance in *cortex vs. ivory* mKO experiments.

| Species | sgRNA(s) | Inj. Embryos Ninj | L1 larvae Nhat | Pupae | Adults Nadu | Crispants Nmut | Hatching Rate Nhat/Ninj | Survival Rate Nadu/Ninj | Hatching Rate Nhat/Ninj | Penetrance Nmut/Ninj |
| --- | --- | --- | --- | --- | --- | --- | --- | --- | --- | --- |
| cortex exon mKOs<br>Livraghi et al. 2016 | *H. erato* | 3122 | 1029 | 511 | 357 | 19 | 33.0% | 11.4% | 0.6% | 5.3% |
|  | *H. charithonia* | 229 | 145 | 61 | 49 | 2 | 63.3% | 21.4% | 0.9% | 4.1% |
|  | *H. melpomene* | 782 | 377 | 195 | 146 | 7 | 48.2% | 18.7% | 0.9% | 4.8% |
| This study | *V. cardui* | 674 | 370 | 53 | 50 | 0 | 54.9% | 7.4% | 0.0% | 0.0% |
|  | **total** | **4807** | **1921** | **820** | **602** | **28** | **40.0%** | **12.5%** | **0.6%** | **4.7%** |
| ivory promoter mKOs | *H. erato* | 294 | 223 | 104 | 74 | 42 | 75.9% | 25.2% | 14.3% | 56.8% |
|  | *H. charithonia* | 917 | 441 | 73 | 47 | 22 | 48.1% | 5.1% | 2.4% | 46.8% |
|  | *V. cardui* | 3100 | 323 | 602 | 331 | 266 | 10.4% | 10.7% | 8.6% | 80.4% |
|  | *A. incarnata* | 74 | 36 | 16 | 13 | 12 | 48.6% | 17.6% | 16.2% | 92.3% |
|  | *D. plexippus* | 100 | 24 | 7 | 2 | 2 | 24.0% | 2.0% | 2.0% | 100.0% |
|  | **total** | **4485** | **1047** | **802** | **467** | **344** | **23.3%** | **10.4%** | **7.7%** | **73.7%** |

**Table S3.**
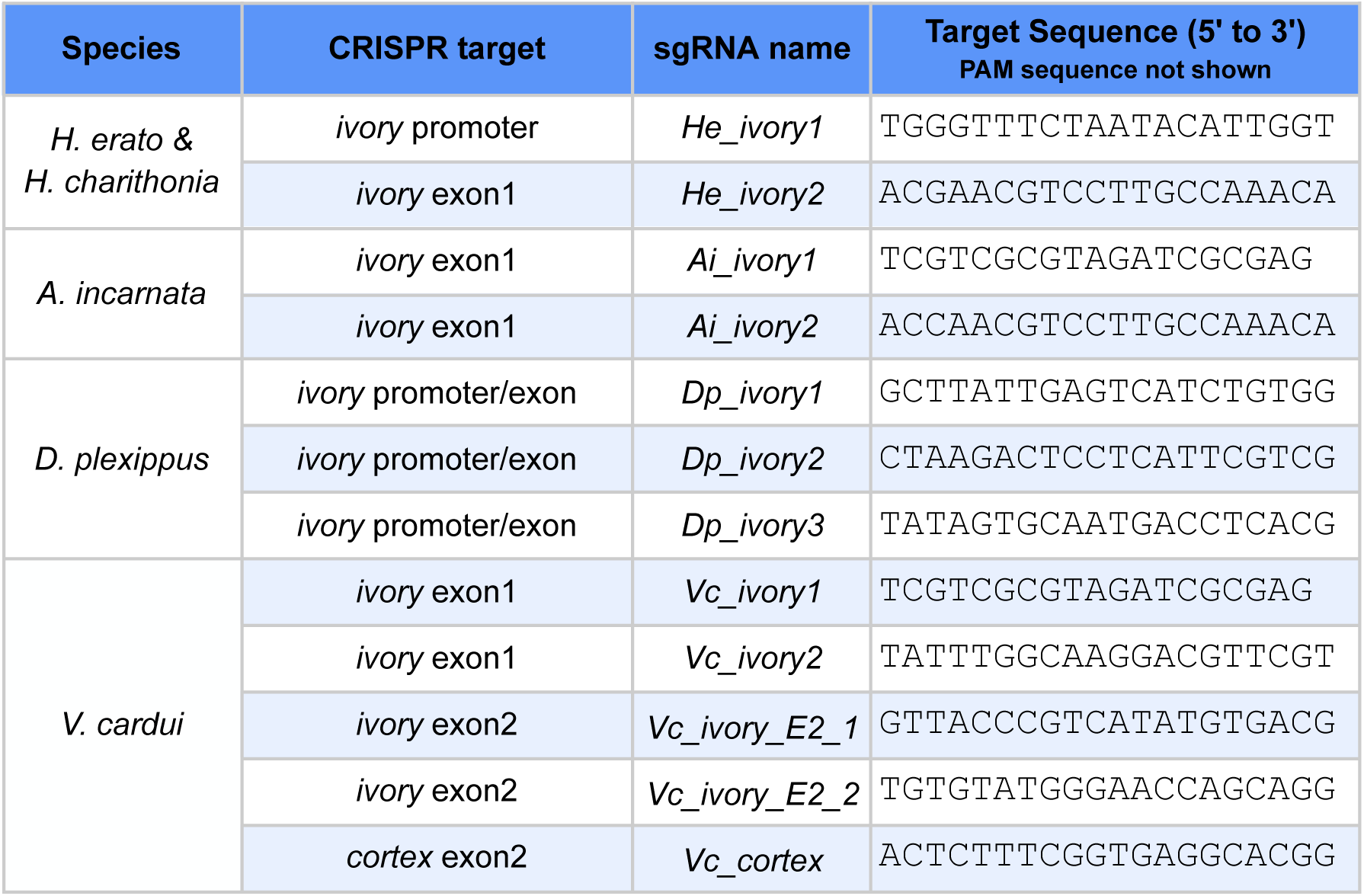
sgRNA targets used in CRISPR mKO experiments.

**Table S4.**
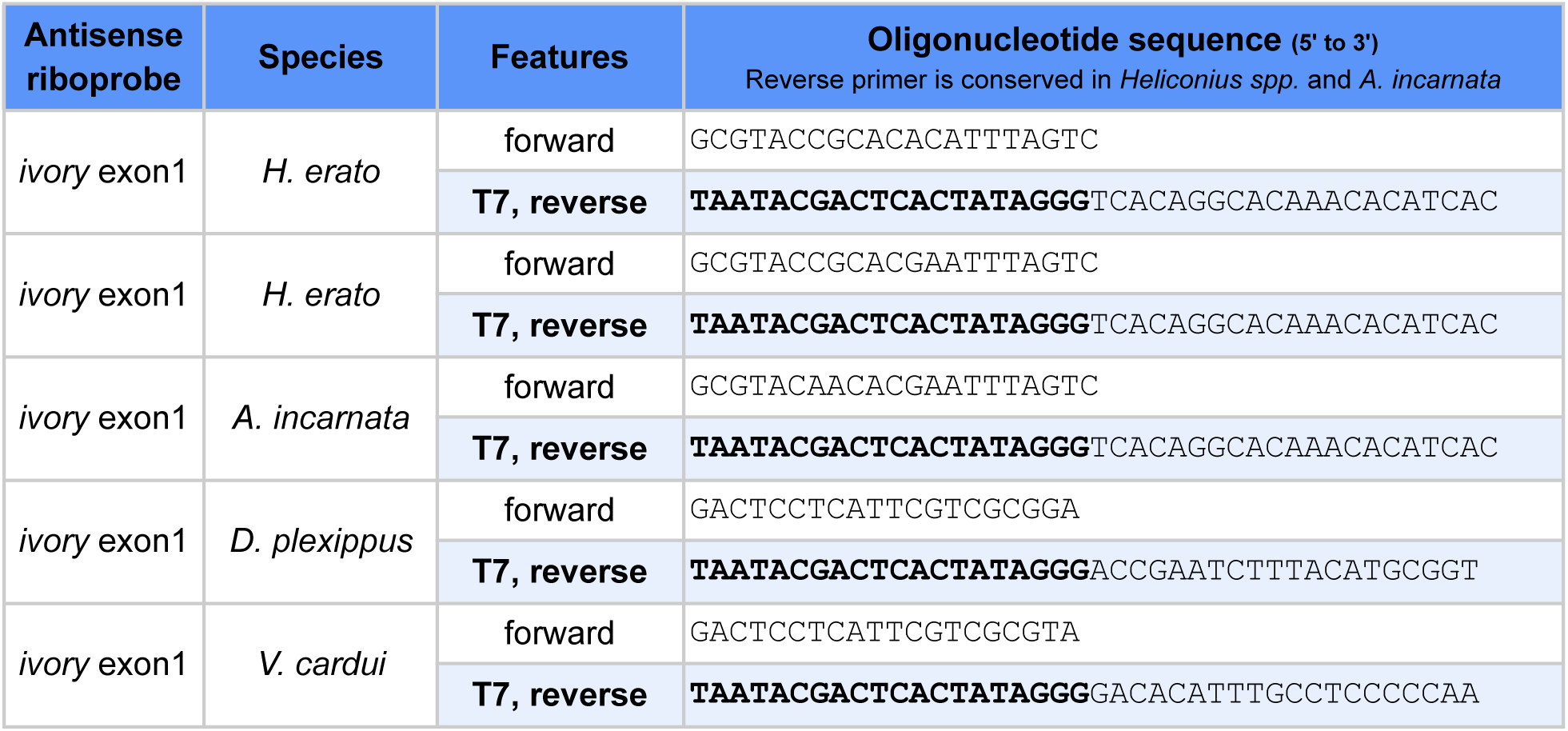
Oligonucleotides used for amplification of ISH riboprobe templates.

**Table S5.**
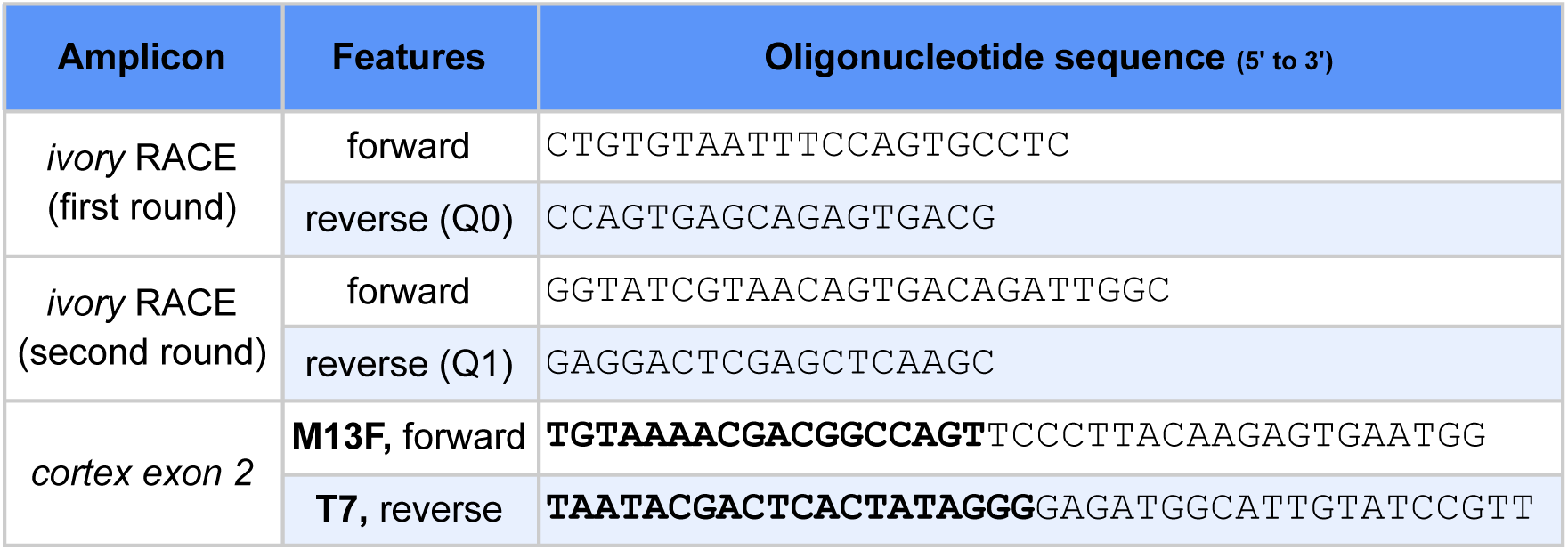
Oligonucleotides used in 3’ RACE and PCR amplification.

## Notes

https://osf.io/Q3SY7/

### Competing Interest Statement

The authors have declared no competing interest.

